# Regulatory T cells establish an IL-10–IL10Rα immunometabolic checkpoint that limits HSL activation and lipolysis

**DOI:** 10.64898/2026.07.12.738050

**Authors:** Ramazan Yildiz, Kajal Davi, Niki F. Brisnovali, James W.R. McMullen, Chung Hwan Cho, Khatanzul Ganbold, YoungUk Jang, Njeri Z.R. Sparman, Aidan Warnock, Gabriel Deards, Leigh Goedeke, Prashant Rajbhandari

## Abstract

Adipose tissue harbors a significant population of regulatory T (Treg) cells that enforce immune homeostasis, yet whether Tregs function as an immunometabolic checkpoint to directly regulate core adipocyte signaling programs remains incompletely defined. Here we show that adipose Tregs function as a dominant, time-dependent checkpoint on β-adrenergic signal-driven lipolytic program and signal transduction in adipocytes. Our integrated scRNA-seq, flow cytometry, and phosphoproteomics data show that prolonged adrenergic stimulation induces a progressive attenuation of activation of key lipase hormone-sensitive lipase (HSL) that coincides with Treg depletion in circulation and accumulation within white adipose tissue. Genetic perturbations establish Treg-derived interleukin-10 (IL-10) as the key mediator of this brake. IL-10 signaling through adipocyte IL-10Rα suppresses adrenergic HSL activation and rewires downstream signaling nodes that govern catecholamine responsiveness, lipolysis, and systemic energy homeostasis. Mechanistically, IL-10Rα engages a STAT3-dependent transcriptional program that induces the G-protein regulators RGS2 and RGS3, diminished PKA flux to HSL that reinforces suppression of the HSL activation state and lipolysis. Together, these findings define an adrenergic–immune feedback circuit in which Tregs fine tune the amplitude and duration of catecholamine responsiveness in adipocytes, establishing immune control of a core lipolytic pathway with implications for obesity-associated adipose dysfunction.

## Introduction

Lipolysis is a central adipocyte function that mobilizes stored triglycerides during fasting, cold exposure and sympathetic activation ^1,2^. Catecholamines drive this response through β-adrenergic activation of the Gs–cAMP–PKA pathway, leading to phosphorylation of hormone-sensitive lipase (HSL) and release of fatty acids and glycerol ^1,2^. Although acute lipolysis is required to meet energetic demand, sustained or excessive lipid mobilization can cause adipose dysfunction, promote ectopic lipid deposition, and contribute to systemic metabolic dysfunction ^3–6^. Thus, adipocytes must not only activate lipolysis rapidly, but also terminate or restrain catecholamine signaling over time ^1,2,7^.

Immune cells are now recognized as important regulators of adipose tissue metabolism, including insulin sensitivity, thermogenesis, inflammation and tissue remodeling ^8–14^. Several immune populations influence adipocyte lipid flux indirectly by shaping inflammatory tone, sympathetic innervation, or catecholamine availability ^1,15–17^. However, most studies of immune control of lipolysis have focused on chronic inflammation, aging, obesity or indirect regulation of adipocyte function, rather than on how adipose tissue actively restrains catecholamine-driven lipolytic tonality ^18^.

Regulatory T cells (Tregs) are among the most highly tissue-adapted immune cells in adipose tissue ^19–21^. Tregs have been linked to improved insulin sensitivity, suppression of adipose inflammation and preservation of tissue function during metabolic stress ^20–24^. In some contexts, Treg depletion or expansion alters systemic lipid handling and adipose tissue remodeling, suggesting that Tregs may influence adipocyte metabolism ^24–27^. Yet whether Tregs directly control the canonical adipocyte lipolytic machinery, and whether Treg-derived factors tune β-adrenergic PKA–HSL signaling, has remained unresolved.

Here we identify a delayed immune feedback circuit that restrains adipocyte catecholamine responsiveness. We show that prolonged β3-adrenergic stimulation first induces a rapid HSL phosphorylation burst and then recruits an IL-10+Treg population to white adipose tissue (WAT). Treg-derived IL-10 acts directly through adipocyte IL10Rα to suppress catecholamine-induced HSL phosphorylation and lipolysis. Mechanistically, IL-10 induces the G protein regulators RGS2 and RGS3, establishing a negative-feedback brake on β-adrenergic signaling upstream of PKA. Loss of Treg-derived IL-10 or adipocyte IL10Rα abolishes this anti-lipolytic program. These findings distinguish Treg control of adipocyte metabolism from classical immune suppression. Rather than simply limiting inflammation, adipose Tregs directly tune adipocyte catecholamine sensitivity through an IL-10–IL10Rα–RGS axis. This defines an immunometabolic checkpoint for lipid mobilization and reveals how immune cells terminate sustained adrenergic lipolysis during metabolic stress.

## Results

### Single cell data show prolonged adrenergic activation elicits a dynamic Treg response in WAT

Since most studies have primarily used short β3-adrenergic agonist CL-316,243 (CL) challenges of 15–30 minutes to capture peak pHSL activation, we aimed to define how WAT integrates lipolytic stimulus over time. We treated C57BL/6 mice with CL and profiled pHSL (pHSL Ser563) in both inguinal (iWAT) and gonadal (gWAT) depots across a 0h, 0.5h, 3h, 8h, and 48 h (injection every 24h) time course. pHSL peaked at 0.5 h after CL injection in both depots and then progressively declined, returning to and dropping below baseline by 48 h (Fig. 1, A and B; quantification by densitometry, P < 0.0001 versus 0.5 h peak; matched unchanged total HSL loading shown in fig. S1A) despite presence of CL. The amplitude of the early pHSL response was comparable between depots, and so was the kinetics of return to baseline between iWAT and gWAT, suggesting that an active suppressive program rather than ligand clearance alone drives termination of catecholamine signaling.

**Fig. 1.**
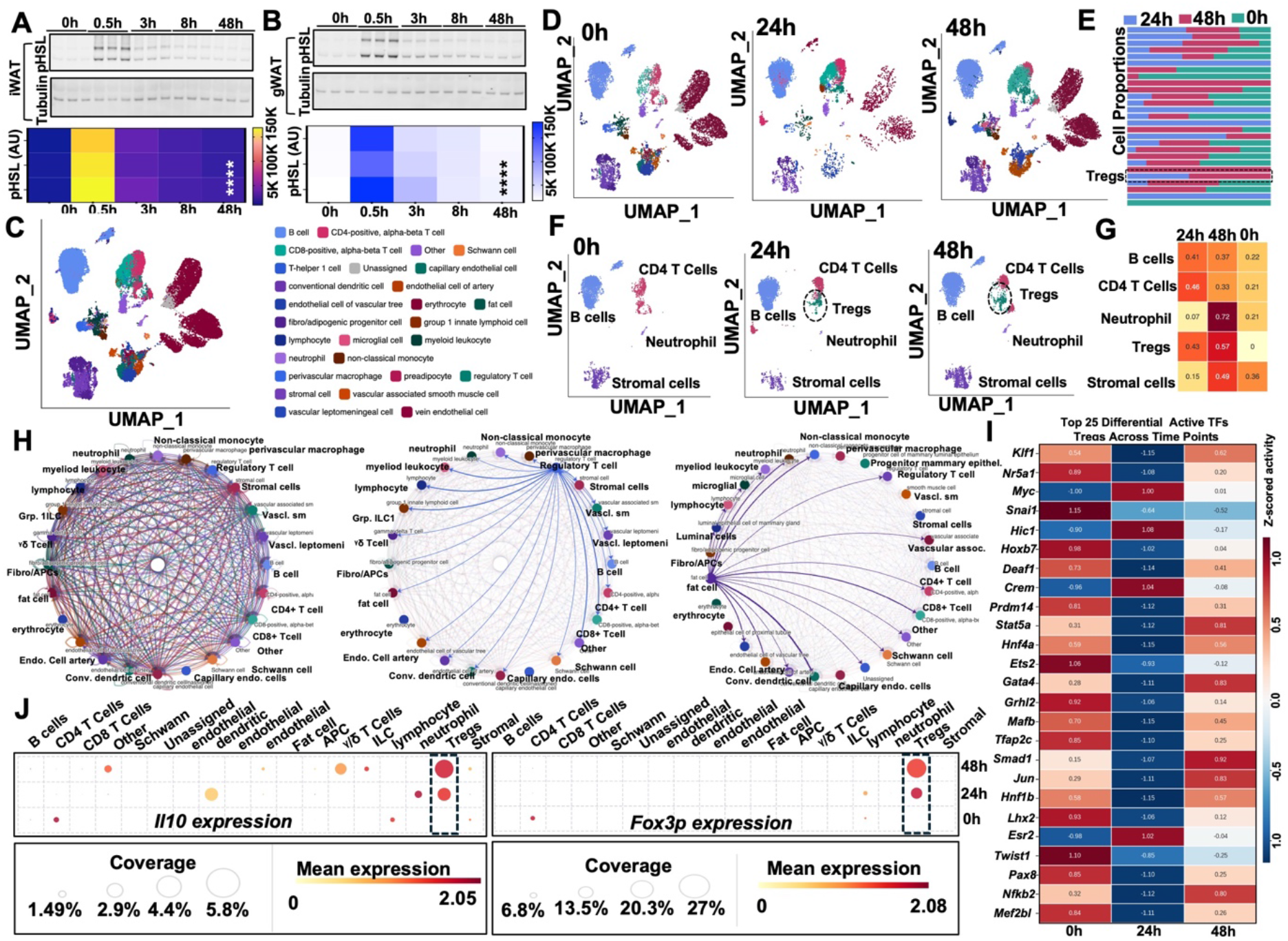
Catecholamine stimulation drives a transient lipolytic burst and a delayed expansion of IL-10–producing Tregs. (A and B) Western blot and densitometric heatmap of phosphorylated hormone-sensitive lipase (pHSL Ser563) in iWAT (A) and gWAT (B) over a 0–48 h time course after intraperitoneal injection of CL-316,243 (1 mg/kg). Tubulin, loading control; n = 3 mice per time point; ****P < 0.0001 versus 0.5 h peak by one-way ANOVA with Tukey post hoc. (C) UMAP embeddings of iWAT stromal-vascular fraction (SVF) scRNA-seq at 0 h, 24 h and 48 h after CL (≥ 38,000 cells across conditions). (D) Composite UMAP across all time points colored by cell-type annotation. (E) Per-cell-type stacked-bar composition across time points highlighting Treg expansion (dashed box). (F) Cell-type-resolved UMAPs across time points highlighting the abundance of a discrete Treg cluster (dashed circles) at 24 and 48 h. (G) Heatmap of per-cell-type proportional changes across 24 h, 48 h and 0 h. (H) CellChat-derived cell–cell communication networks across 0 h, 24 h and 48 h, demonstrating reorganization toward Treg- and adipocyte-centered outgoing signaling. (I) Heatmap of the top 25 differentially active transcription factors (SCENIC) in Tregs across time points (z-scored regulon activity). (J) Dot plots of Il10 (left) and Foxp3 (right) expression across cell types and time points (dot size, fraction of cells expressing; color, mean expression). Dashed boxes highlight Tregs as the dominant source of Il10 after CL stimulation.

To identify the cellular source of this restraint, we performed single-cell RNA sequencing (scRNA-seq) of the iWAT stromal-vascular fraction (SVF) at 0, 24 and 48 h after CL administration (Fig. 1C). Among identified cell populations, the FOXP3+ regulatory T cell (Treg) compartment showed the most pronounced temporal change: Tregs were rare at 0 h, expanded markedly by 24 h, and remained elevated at 48 h (Fig. 1D,E). Visualizing this dynamic in lineage-resolved UMAPs confirmed the appearance of a discrete Treg cluster at 24–48 h (Fig. 1F). Per-cell-type proportion analysis identified Tregs as the cell type with the largest fold-change between 24 h and 48 h, coincident with the drastic drop in HSL activation (Fig. 1G). Catecholamine treatment also reorganized predicted cell–cell communication, with focused Tregs and adipocytes acquiring broad connectivity at 24–48 h (Fig. 1H). We replicated these findings independently using data analysis and consensus cell annotation atlas for mouse adipose tissue (Fig. S1B) ^28^. We further used our published dataset ^10^ from mice treated with CL for 4 days and comparing CL-treated and saline-treated iWAT SVF directly, the frequency of IL-10–expressing Tregs rose 2.28-fold (CL: 6.89% versus saline: 3.02%; Wilcoxon P = 0.0048), and mean IL-10 expression per Treg rose 2.53-fold (fig. S1C-F).

We tested whether expanding Tregs were transcriptionally programmed to produce IL-10, we computed single-cell transcription-factor activity scores (SCENIC) ^29^. Tregs at 24–48 h displayed a coordinated increase in the activity of regulators governing Treg effector function and IL-10 transcription, including Stat5a, Hnf1b, Smad1, Crem, Hic1, Nr5a1 and Mafb (Fig. 1I). Consistent with this rewiring, *Il10* transcripts were almost undetectable in baseline iWAT but became selectively enriched in the Treg cluster at 24 h and 48 h, with concomitant induction of Foxp3 (Fig. 1J). Tregs were the dominant cellular source of *Il10* in CL-stimulated WAT, surpassing macrophages, conventional dendritic cells, B cells and innate lymphoid cells by both fraction expressing and mean expression.

The cell-type-resolved chemokine landscape further showed that CL-activated macrophages, dendritic cells and stromal APCs up-regulated the canonical Treg-recruiting chemokines Ccl2, Ccl1, Ccl17, Ccl22 and Cxcl12 by 24 h (Fig. S1, G and H) ^30^. Together, these data establish a previously unknown kinetic framework in which catecholamine signaling first triggers a lipolytic burst and then mobilizes an IL-10+Treg population that is temporally positioned to dampen the adrenergic tonality and HSL activation.

### IL-10–producing Tregs directly acts on adipocytes in vitro and in vivo to restrict HSL activation

To further validate our transcriptomics data, we performed a multiparameter flow cytometry on SVFs of both gWAT and iWAT and on peripheral blood from C57BL/6J mice harvested before treatment (PBS) (n=6) and 0.5 h (n=6) or 48 h (n=6) after a single (0.5h) or two 24h doses (48h) of CL. First, live singlets were sequentially gated on FSC-H/CD45+ for total leukocytes, CD3+CD4+ for conventional CD4+ T cells, CD25+FoxP3+ for regulatory T cells and IL10+ for IL-10-producing/expressing regulatory T cells (Fig. 2A; Fig. S2A-H). In agreement with our scRNA-seq data, we noticed a marked accumulation of IL-10+Tregs after 48H CL treatment (Fig. 2A). We then performed a comprehensive immune profiling in WAT (see Methods), and at 0.5 h, coincident with the peak of catecholamine-driven HSL phosphorylation, adipose-resident leukocytes were uniformly depleted: gWAT CD45^+^ leukocytes (log2FC = −1.1), CD4^+^ T cells (−2.3, P < 0.05), Foxp3^+^ Tregs (−3.9, P < 0.05) and IL-10^+^ Tregs (−4.2, P < 0.05) all dropped relative to PBS, mirrored by a transient decline in circulating leukocytes, B220^+^ B cells, CD3^+^CD90^+^ T cells, and CD4^+^ T cells in plasma (Fig. 2B; Fig. S2, C and F). By 48 h, in gWAT total leukocytes expanded 6.5-fold (log2FC = +2.7, P < 0.05) and CD4^+^ T cells, Foxp3^+^ Tregs and IL-10^+^ Tregs each rose ∼5-fold (log2FC = +2.6, +2.3, and +2.3 respectively; all p< 0.05; Fig. 2B). Absolute densities confirmed the magnitude, gWAT IL-10^+^ Foxp3^+^ Tregs rose from ∼25 to ∼120 cells/mg (P < 0.01; fig. S2B). The immune leukocyte reorganization was also seen in iWAT, which showed induction of CD4^+^ T cells (log2FC = +1.7, p< 0.05) and Foxp3^+^ Tregs (+1.1, p< 0.05) and IL-10^+^ Treg accumulation (Fig. S2D), and blood IL-10^+^ Tregs remained low (−1.3, P < 0.05, Fig. S2E). An elaborate multiple timepoints (Fig. S2F-H, n=3/condition) resolved the dynamics where blood IL-10^+^ Tregs decreased from ∼1,600 to ∼250 cells/ml within the first 8 h and remained suppressed for 48 h, while iWAT and gWAT IL-10^+^ Treg frequencies markedly increased by 48h, with IL-10^+^ Tregs reaching ∼8% of CD4^+^ T cells by 48 h. Body, iWAT and gWAT weights all decreased significantly between PBS and CL 48 h, with the largest absolute mass loss in gWAT (Fig. S2A). Together these data establish that the IL-10–producing Treg pool is low during the immediate-early phase of catecholamine action and *redistributes* from circulation to WAT on a 24–48 h timescale, positioning adipose IL-10^+^ Tregs as a candidate paracrine source of a delayed brake on HSL activation and lipolysis.

**Fig. 2.**
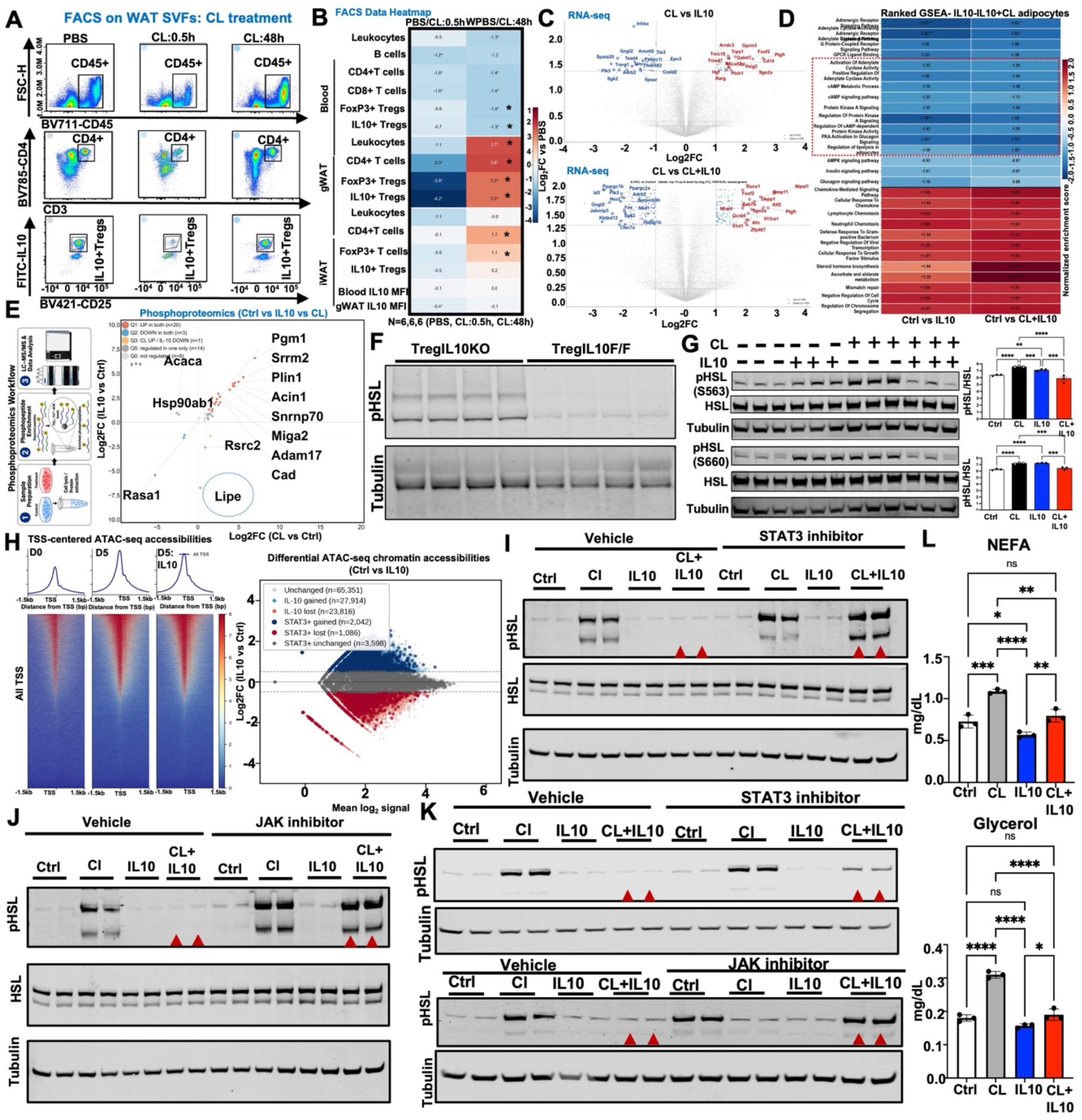
Treg-derived IL-10 acts on adipocytes to suppress catecholamine-driven HSL phosphorylation. (A) Representative FACS gating cascade on stromal-vascular fractions (SVFs) of gWAT from C57BL/6J mice treated with PBS, CL-316,243 (CL) for 0.5 h, or CL for 48 h: events were gated on FSC-H × BV711–CD45 (top row, CD45+), CD3 × BV785–CD4 (middle row, CD3+CD4+), and BV421–CD25 × FITC–IL-10 (bottom row, IL-10+ Tregs within CD4+CD25+Foxp3+). (B) Heatmap of log2 fold-change versus PBS for the indicated immune populations — leukocytes, B cells, CD4+ T cells, CD8+ T cells, Foxp3+ Tregs, IL-10+ Tregs — and IL-10 mean fluorescence intensity (MFI) in blood, gWAT and iWAT at CL 0.5 h and CL 48 h. Color scale: log2FC vs PBS (−4 to +4). N = 6 mice per condition. P < 0.05 vs PBS by two-tailed Student’s t-test. (C) Volcano plots of bulk RNA-seq in 10T1/2 differentiated adipocytes treated with CL or IL-10 (top) or CL or CL+IL-10 (bottom) versus control; named genes are the top 15 up- and down-regulated transcripts (|log2FC| > 1, FDR < 0.05). (D) Heatmap of normalized enrichment scores (NES) for selected MSigDB GO/Reactome/KEGG gene sets from ranked GSEA of the comparisons in C; values shown only for FDR < 0.05. The boxed cluster (red dashed outline) highlights the coordinated IL-10-mediated suppression of the adrenergic/cAMP/PKA/lipolysis axis (NES −1.08 to −1.85). (E) Workflow and scatter plot of quantitative phosphoproteomics (LC-MS/MS, n = 3 per group) comparing IL-10/Ctrl (y-axis) versus CL/Ctrl (x-axis) phosphosite log2FC. Quadrant colors: Q1 up in both (n = 20), Q2 down in both (n = 3), Q3 CL-up/IL-10-down (n = 1), Q4 (n = 0), Q5 regulated in one only (n = 14), Q0 not regulated (n = 6). Labeled proteins include the HSL ortholog Lipe (encircled) and lipid-droplet/lipolysis-proximal substrates (Plin1, Acaca, Cad, Adam17, Miga2, Pgm1, Srrm2, Acin1, Snrnp70, Rasa1, Hsp90ab1). (F) Western blot of pHSL and tubulin in gWAT from TregIL10KO and TregIL10F/F littermate controls after CL stimulation; four biological replicates per genotype. (G) Representative western blots and densitometric quantification (right) of pHSL Ser563 and Ser660, total HSL and tubulin in adipocytes treated with Ctrl, CL, IL-10 or CL+IL-10. n = 3 per group; mean ± s.e.m.; P < 0.01, P < 0.001, P < 0.0001 by one-way ANOVA with Tukey’s post-hoc test. (H) Left: TSS-centered ATAC-seq accessibility aggregate profiles and heatmaps at D0, D5, and D5+IL-10 in primary adipocytes (color scale, normalized signal). Right: differential ATAC-seq peak scatter (Ctrl vs IL-10) annotated by TOBIAS/BINDetect STAT3 footprint status, unchanged (n = 65,351), IL-10 gained (n = 27,914), IL-10 lost (n = 23,816), STAT3-footprinted gained (n = 2,042), STAT3-footprinted lost (n = 1,086), STAT3-footprinted unchanged (n = 3,598). (I) Western blot of pHSL, total HSL and tubulin in human primary adipocytes treated with Ctrl, CL, IL-10 or CL+IL-10 in the presence of vehicle (left) or a STAT3 inhibitor (right). Red arrowheads show IL-10-suppressed pHSL bands restored by STAT3 inhibition. (J) Western blot of pHSL, total HSL and tubulin in human primary adipocytes treated with Ctrl, CL, IL-10 or CL+IL-10 in the presence of vehicle or a JAK inhibitor; red arrowheads, IL-10-suppressed bands restored by JAK inhibition. (K) Independent mouse adipocyte replicates of the STAT3-inhibitor (top) and JAK-inhibitor (bottom) rescue experiments with parallel pHSL/tubulin blots; red arrowheads as in I and J. (L) Serum NEFA (Up) and glycerol (down) across Ctrl, CL, IL-10 and CL+IL-10 (n = 3 per group). P < 0.05, *P < 0.01, P < 0.001, ***P < 0.0001 by one-way ANOVA with Tukey’s; n.s., not significant.

IL-10Rα is expressed in mouse and adipocytes ^9–11^ (Fig. S2I) and to test whether IL-10 directly modifies the adipocyte response to CL, we differentiated preadipocytes and treated them with vehicle, CL, IL-10, or CL+IL-10 for 6 h, followed by bulk RNA-seq, quantitative phosphoproteomics, and western blotting. Volcano analysis (|log2FC| > 1, FDR < 0.05) identified ∼190 differentially expressed genes in the CL vs IL-10 comparison and a markedly larger ∼280-gene response in the CL vs CL+IL-10 comparison, with co-treated cells acquiring distinct up-regulated transcripts (*Runx1*, *Mmp11*, *Nipal1*, *Klf2*, *Ahr*, *Il13ra1*, *Zfp467, Ptgfr*) and suppressing CL-induced *Ppargc1a/b*, *Adrb2*, *Sgk2*, *Jakmip3* and *Pla2g1b* (Fig. 2C). Ranked GSEA on curated MSigDB collections (GO/Reactome/KEGG, FDR < 0.05) revealed that IL-10 — alone and in combination with CL — selectively down-regulated the pro-lipolytic axis: adrenergic receptor signaling (NES = −1.57 / −1.63), adenylate cyclase activation (−1.85 / −1.66), cAMP metabolic process (−1.30 / −1.38), cAMP signaling (−1.22 / −1.13), PKA signaling (−1.25 / −1.42), and regulation of lipolysis in adipocytes (−1.08 / −1.43), with concomitant up-regulation of chemokine-mediated signaling and immune-cell chemotaxis (NES = +1.58 to +1.83) (Fig. 2D, boxed cluster). Quantitative phosphoproteomics (LC-MS/MS, n = 3 combined per group) on IL-10/Ctrl versus CL/Ctrl identified 44 concordantly regulated phosphosites: 20 up in both (Q1), 3 down in both (Q2), 1 CL-up/IL10-down (Q3), and 14 regulated in only one condition (Q5; Fig. 2E). Among the directly affected substrates were lipid-droplet– and HSL-proximal proteins, Lipe (HSL, encircled), Plin1, Acaca, Cad, Adam17, Miga2, Pgm1, Srrm2, Acin1, Snrnp70 and Rasa1, establishing that IL-10 remodels the catecholamine-driven phosphoproteome at the level of the lipolytic machinery. Direct validation in vivo by western blot confirmed our findings, where Treg-specific IL-10–knockout mice (TregIL10KO, *Foxp3*-Cre × *Il10*^fl/fl^), gWAT and iWAT pHSL was markedly elevated relative to TregIL10F/F littermate controls (Fig. 2F and Fig. S2J,K), and in ex vivo differentiated adipocytes co-treatment with IL-10 attenuated CL-induced phosphorylation at both regulatory sites of HSL, Ser^563^ (pHSL/HSL drop ∼65%) and Ser^660^ (∼60%) with ****p < 0.0001 (Ctrl/CL/IL10/CL+IL10, one-way ANOVA with Tukey’s; Fig. 2G, S2L). IL-10 did not alter total HSL protein, indicating regulation at the phosphorylation step rather than at protein abundance. Bioenergetic analysis by Seahorse Mitochondrial Stress test confirmed that the CL-induced increase in oxygen consumption rates (OCR) was abolished by IL-10 co-treatment (CL+IL-10 OCR was ∼50% lower than CL alone; *P < 0.05, P < 0.01;* Fig. S2M), consistent with reduced fatty-acid β-oxidation.

To identify the transcriptional mechanism underlying the IL-10 response, we performed ATAC-seq on primary adipocytes at D0, D5, and D5+IL-10. TSS-centered accessibility heatmaps and aggregate profiles showed that IL-10 broadly opens chromatin at adipocyte gene loci and promoter regions during differentiation (Fig. 2H, left). Genome-wide differential peak calling identified 27,914 IL-10–gained, 23,816 IL-10–lost, and 65,351 unchanged peaks (Fig. 2H, right). To assign these accessibility changes to specific transcription factors, we performed BINDetect footprinting ^31^ at TOBIAS-annotated STAT3 motifs and overlaid the calls on the IL-10–differential peak set. STAT3 footprints were significantly enriched among IL-10–gained peaks (7.3% vs 5.5% in unchanged peaks and 4.6% in IL-10–lost peaks; *P* = 6.0 × 10^−40^; Fig. S2N), and 2,042 IL-10–gained peaks and 1,086 IL-10–lost peaks carried high-confidence STAT3 footprints (Fig. 2H, right). Independent HOMER known-motif enrichment on IL-10–differential peaks (n = 221 lost; n = 101 gained) recovered Atf4, Chop, CEBP:AP1, MRE, CEBP:CEBP, GRE, PGR, ARE, AARE and PR motifs among IL-10–lost peaks and ZBTB18, HLF, TCF4, NeuroG2, Twist2, BHLHA15, Olig2, Atoh1, AT1G49560, NFIL3 motifs among IL-10–gained peaks (Fig. S2O), consistent with a STAT3-mediated remodeling that selectively closes adrenergic stress-response chromatin while opening adipocyte/lineage-restricted enhancers. Our functional experiments confirmed STAT3 as the relevant IL-10 transducer, small-molecule STAT3 inhibitor rescued CL-driven HSL phosphorylation in CL+IL-10–treated human and mouse adipocytes (Fig. 2I,K, red arrowheads), and a JAK inhibitor, upstream of STAT3, showed the same rescue of CL-driven HSL phosphorylation in human and mouse differentiated adipocytes (Fig. 2J,K), without altering vehicle pHSL. Together, these data confirm our scRNA-seq data and identify Treg-derived IL-10 as a brake on HSL activation and β-adrenergic lipolytic signaling across species.

To determine whether the molecular brake translates into reduced lipolytic flux, we measured serum non-esterified fatty acids (NEFA) and glycerol, the two principal hydrolysis products of HSL/ATGL action, across treatment groups. CL elevated NEFA from 0.72 to 1.08 mg/dL (P < 0.001), and co-treatment with IL-10 reduced NEFA back to 0.78 mg/dL (P < 0.01 vs CL); IL-10 alone slightly lowered baseline NEFA (0.56 mg/dL, P < 0.0001 vs CL) (Fig. 2L). Glycerol release showed the same pattern (CL: 0.31 vs CL+IL-10: 0.19 mg/dL; P < 0.05; Fig. 2J). Untargeted lipidomics of treated adipocytes resolved this functional brake at the molecular-species level. Principal-component analysis separated Ctrl, CL, and IL-10+CL on PC1 (79.4%) and PC2 (20.6%; Fig. S2P). Plotting the IL-10 effect (log2FC IL-10+CL/CL) against the CL effect (log2FC CL/Ctrl) defined two reciprocal rescue features: a TG/DG rescue region (CL↓ / IL-10↑) and an FA rescue region (CL↑ / IL-10↓) (Fig. S2Q). At the lipid class level, IL-10 attenuated the CL-driven depletion of triglycerides (TG), diacylglycerols (DG), phosphatidylethanolamines (PE), cholesteryl esters (CE) and phosphatidic acid (PA), and reversed the CL-induced rise in free fatty acids (FA) (Fig. S2R,S). At the species level, this was resolved in the per-species z-score heatmap (Fig. S2R) and in absolute abundance plots: TG 60:12-FA22:6 (0.56 Ctrl / 0.01 CL / 0.18 IL-10+CL) and DG 22:0_20:0 (0.54 / 0.01 / 0.17) were rescued by IL-10, while free FAs FA 22:4 (3.60 / 4.03 / 3.82) and FA 20:4 (6.31 / 6.40 / 5.95) were partly rescued (Fig. S2T,U). Across diverse experimental modalities, transcriptome, phosphoproteome, chromatin, oxygen consumption, immunoblot, mouse model, and the lipidome, IL-10 acts on adipocytes as a brake on β-adrenergic lipolysis and HSL activation.

### Adipocyte IL-10Rα is required in vivo to restrict adipocyte HSL activation and lipolysis

Next, we tested the requirement for adipocyte-intrinsic IL-10 action in vivo (Fig. 3A). Since we noticed that IL10 also blocks HSL activation in human adipocytes, we first asked whether this axis is functional in humans. Large-scale multi-ancestry lipids GWAS meta-analysis (METSIM, FINRISK and HCHS/SOL) ^32–35^ showed that common non-coding variants at the IL10RA locus on chromosome 11 (rs74737837-G, rs36119055-T) were associated with circulating triglycerides at genome-wide significance in European-ancestry (P = 8e^-1^) and Hispanic/Latino (P = 1e^-13^) cohorts (Fig. 3B). In adipose-biopsy/clinical-trait data ^36^, IL-10RA expression in WAT correlated positively with waist-hip ratio (WHR), waist, HOMA-IR, BMI, circulating insulin, circulating glucose, HbA1c, circulating CRP, circulating cholesterol, circulating LDL, fat-cell volume and negatively with circulating-HDL, basal/isoproterenol-stimulated lipolytic ratio (Fig. 3C), supporting an adipocyte-autonomous, physiologically relevant role for IL-10R in human adipocyte lipolysis.

**Fig. 3.**
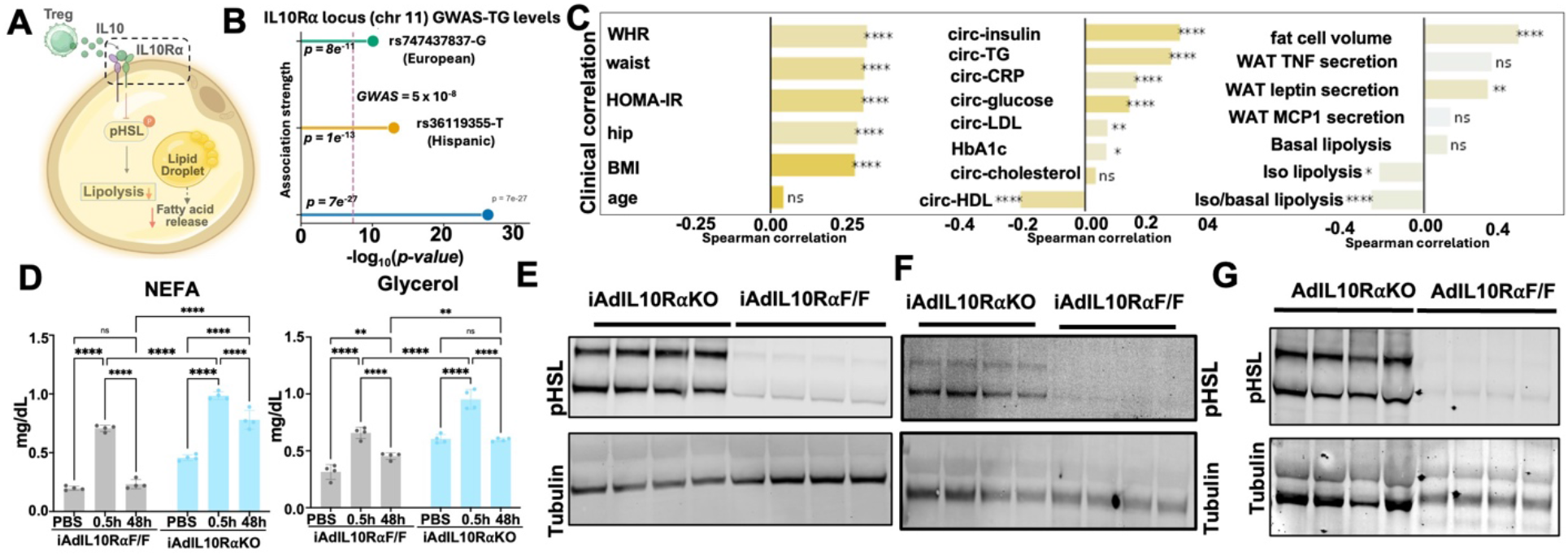
Adipocyte IL-10Rα is required in vivo to limit adipocyte lipolysis and HSL activation. (A) Schematic of IL-10–IL-10Rα–pHSL signaling in the adipocyte (Treg IL-10 to IL-10Rα to suppression of pHSL and lipolysis). (B) Manhattan-style plot of the IL10RA locus (chromosome 11) showing GWAS associations with circulating triglycerides in European and Hispanic/Latino cohorts. (C) Forest plots of Spearman correlations between IL10RA expression in WAT and metabolic/clinical traits (left and middle) and adipose secretion / lipolysis readouts (right). (D) Circulating NEFA and glycerol across PBS, CL 0.5 h and 48 h in iAdIL10RαF/F and iAdIL10RαKO. (E and F) pHSL western blots in iWAT (E) and gWAT (F) of iAdIL10RαKO versus iAdIL10RαF/F mice across 48h CL time points. (G) pHSL western blot in AdIL10RαF/F and AdIL10RαKO WAT. (Mean ± SEM; \**p < 0.01, p < 0.001,* ***p < 0.0001. Panel A was made using BioRender.

We generated an inducible, adipocyte-specific IL-10Rα knockout (iAdIL10RαKO; *Adipoq-rtTA* × *TRE-Cre* × *Il10rafl/fl* + doxycycline) (Fig. 3D). Compared to iAdIL10RαF/F littermates, iAdIL10RαKO mice showed persistent elevation of pHSL and not total HSL across the CL time course (48h) in both iWAT and gWAT (Fig. 3, E,F and Fig. S3A,B), with correspondingly elevated circulating NEFA and glycerol (Fig. 3H, ****P < 0.0001). Independent loss-of-function mouse models with constitutive adipocyte IL-10Rα loss (AdIL10RαKO) recapitulated the phenotype (Fig. 3G). The convergence of orthogonal loss-of-function approaches; inducible and constitutive genetic deletion, establishes that adipocyte IL-10Rα is required to set the steady-state setpoint of catecholamine-driven lipolysis, HSL activation, and downstream metabolic phenotype in vivo.

### IL-10 induces the GAPs RGS3 and RGS3 in adipocytes, which is necessary and sufficient to terminate catecholamine signal and HSL activation

To identify the molecular effector that couples adipocyte STAT3 activation to suppression of PKA-driven HSL phosphorylation, we re-examined the IL-10-regulated cAMP/PKA-pathway gene set from Fig. 2D. Among the consistently induced transcripts across IL-10, CL and CL+IL-10, *Rgs3* and *Rgs2*, two regulators of G-protein signaling (RGS) that accelerate GTP hydrolysis on G subunits and terminate adenylate-cyclase activation ^37–43^, emerged as the strongest candidates by both fold-change and reproducibility (Fig. 4A, Fig. S4A-C). IL-10 stimulation alone was sufficient to induce RGS3 protein in adipocytes (Fig. 4B, Fig. S4J) and specificity tests on a panel of co-regulated transcripts IL13RA1, INHBB, and PTGFR (Fig. S4A-C) showed that none were induced at protein level narrowing the relevant effector layer to the RGS family. RGS3 induction required intact adipocyte IL-10 signaling where RGS3 protein was significantly lower in iAdIL10RαKO WAT than in iAdIL10RαF/F adipocytes (Fig. 4D, P = 0.0064) and was reduced in the WATs of TregIL10KO mice compared to controls (Fig. 4C, P = 0.0157). Concordantly, RGS3 was reduced after IL10Rα ASO treatment in vivo (Fig. S4G). Genome-browser analysis of ATAC consensus peaks at the *Rgs3* and *Rgs2* loci confirmed IL-10-gained accessibility flanking the transcription-start sites, with TOBIAS BINDetect calling reproducible STAT3 footprints exclusively in the IL-10 condition (+1.5 kb and +11.5 kb of Rgs3 TSS, FP logFC ≈ +0.20 to +0.83; +72.7 kb of Rgs2 TSS; fig. S4D,E), implicating Rgs2/Rgs3 as direct IL-10-STAT3 targets.

**Fig. 4.**
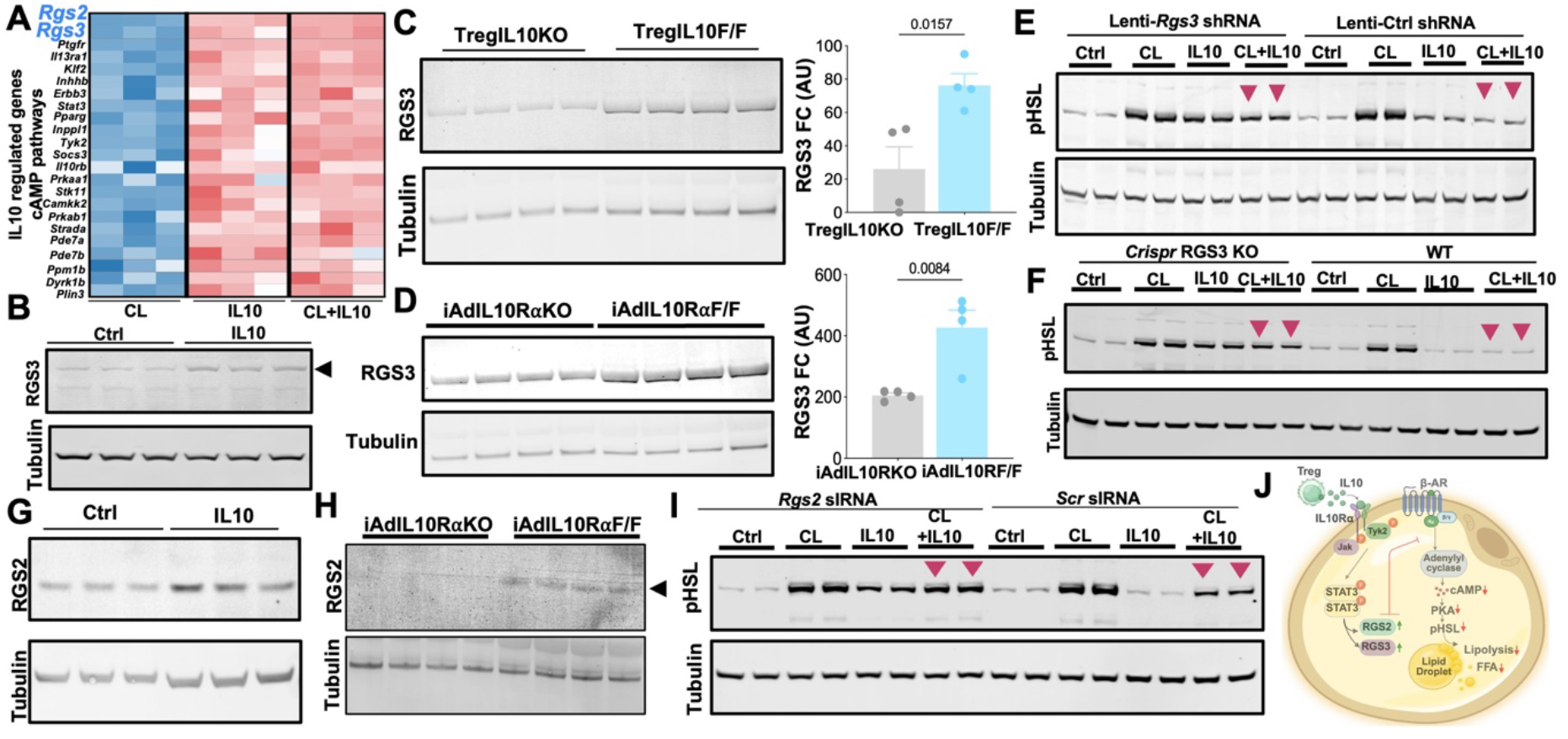
IL-10–STAT3 induces the GAPs RGS3 and RGS2 in adipocytes, which is required for IL-10-mediated suppression of catecholamine-driven HSL phosphorylation. (A) Heatmap of IL-10-regulated cAMP/PKA-pathway genes from Fig. 2C with Rgs2 and Rgs3 highlighted (blue). (B) Representative western blot showing IL-10 induction of RGS3 in differentiated adipocytes. (C and D) RGS3 western blots and densitometric quantification in TregIL10KO versus TregIL10F/F (C; P = 0.0157) and iAdIL10RαKO versus iAdIL10RαF/F (D; P = 0.0064) WAT (n = 4–5 mice per group). (E) pHSL western blots in Lenti-Rgs3 shRNA versus Lenti-Ctrl shRNA adipocytes under Ctrl, CL, IL-10 and CL+IL-10; red arrowheads indicate loss of IL-10-mediated pHSL suppression in shRgs3 cells. (F) pHSL western blots in CRISPR Rgs3-KO versus WT adipocytes across the same four conditions; red arrowheads indicate near complete loss of IL-10-mediated pHSL suppression in RGS3KO cells (G) RGS2 induction by IL-10 in adipocytes. (H) RGS2 western blot in iAdIL10RαKO versus iAdIL10RαF/F WAT. (I) pHSL western blots under Rgs2 siRNA versus scrambled siRNA across the four conditions; red arrowheads indicate loss of IL-10-mediated pHSL suppression in Rgs2 siRNA cells. (J) Cartoon model (made using BioRender) summarizing the Treg-adipocyte axis to dampen lipolysis.

To test sufficiency and necessity, we performed three independent orthogonal loss-of-function experiment: lentiviral Rgs3 shRNA and Scr-control shRNA in differentiated adipocytes (Fig. 4E, Lenti-Rgs3 shRNA reduced RGS3 protein by ≥85%, Fig. S4K, P = 0.0002), siRNA knockdown of Rgs3 (Fig. S4H,I), and CRISPR-Cas9 Rgs3 knockout (Fig. 4F, Fig. S4L). In all the models, IL-10 markedly lost the ability to suppress CL-induced pHSL: Lenti-Rgs3 shRNA cells in the CL+IL-10 condition showed pHSL levels indistinguishable from CL alone, while Lenti-Ctrl shRNA cells retained full IL-10 suppression (Fig. 4E, maroon arrowheads), and the same loss of suppression was even higher in CRISPR *Rgs3* KO adipocytes (Fig. 4F, marron arrowheads). siRNA knockdown of Rgs3 in an independent system reproduced this loss of IL-10-mediated pHSL restraint (Fig. S4H,I), quantitatively defining RGS3 as the dominant effector of IL-10-mediated catecholamine restraint.

We next tested the role of RGS2 in IL-10-mediated inhibition of HSL phosphorylation. In matched human adipose biopsies ^36^, similar to IL-10Rα, RGS2 expression correlated positively with HOMA-IR, WHR, waist, BMI, circulating insulin, glucose and TG, and inversely with iso/basal-stimulated lipolytic ratio (Fig. S4F), consistent with the mouse genetic evidence that the IL-10Rα–STAT3–RGS axis tunes adipocyte catecholamine sensitivity in humans. Similar to RGS3, IL-10 stimulation alone was sufficient to induce RGS2 protein in adipocytes (Fig. 4G, Fig. S4M, p=0.037). RGS2 induction required intact adipocyte IL-10 signaling: RGS3 protein was markedly lower in iAdIL10RαKO WAT than in iAdIL10RαF/F adipocytes (Fig. 4H) and was reduced in the WATs of TregIL10KO mice compared to controls (Fig. S4N). IL-10 lost the ability to suppress CL-induced pHSL in siRNA knockdown of Rgs2 in adipocytes, positioning RGS2 as another effector of IL10 action (Fig. 4I).

Together, these results define a complete immune–adipocyte feedback circuit in which sustained catecholamine stimulation recruits IL-10–competent adipose Tregs, Treg-derived IL-10 activates adipocyte IL10Rα–STAT3 signaling to remodel chromatin and induce Rgs2 and Rgs3, and adipocyte IL10Rα is required in vivo to restrain HSL activation and maintain catecholamine-responsive lipolytic homeostasis. Mechanistically, RGS2 and RGS3 emerge as key downstream effectors of this axis, acting as dominant brakes that terminate cAMP–PKA–HSL signaling. Consistent with this model, loss of Treg-derived IL-10 or adipocyte IL10Rα increases PKA substrate activation in adipocytes relative to controls, supporting a role for the Treg–IL-10–IL10Rα–RGS pathway in limiting adrenergic signal propagation (Fig. 4J and Fig. S4P,Q).

## Discussion

Lipolysis in WAT controls systemic fatty-acid availability and provides lipid substrates for metabolic adaptation during fasting, cold exposure and sympathetic activation. The canonical β-adrenergic–cAMP–PKA–HSL pathway has generally been viewed as an adipocyte-intrinsic, hormone-gated module. Here we identify an additional layer of control: a delayed immune-derived brake in which IL-10-producing regulatory T cells act through adipocyte IL10Rα–STAT3 signaling to restrain HSL activation and lipolysis. This circuit reframes catecholamine-driven lipolysis as a tissue-level program that is initiated by adipocytes but terminated, at least in part, by an inducible immunometabolic checkpoint.

A defining feature of this pathway is its temporal tissue organization. Acute β3-adrenergic stimulation induced a rapid pHSL burst, whereas Treg recruitment/expansion and IL-10 induction emerged at a prolonged duration coinciding with the resolution phase of the lipolytic response. This delay distinguishes the IL-10–IL10Rα axis from intracellular regulators of lipolysis, such as phosphodiesterases, perilipin remodeling, ATGL inhibition, or immediate feedback within the cAMP pathway^1^. Instead, adipose tissue appears to mount a secondary immunoregulatory program that limits the duration of catecholamine responsiveness. This temporal separation may allow WAT to preserve lipid mobilization while preventing sustained or excessive lipolytic flux. The specificity of the circuit is supported by perturbations at multiple levels. Loss of Treg-derived IL-10, inducible and constitutive deletion of adipocyte IL-10Rα, and pharmacological blockade of JAK–STAT3 signaling each impaired the ability of adipose tissue to suppress CL-induced HSL phosphorylation. These observations place Treg-derived IL-10 upstream of an adipocyte-intrinsic receptor pathway and suggest a unique model of suppression of adrenergic-induced WAT inflammation ^44,45^. Hence, Treg-derived IL-10 acts directly on adipocytes to tune the β-adrenergic response. Mechanistically, our data identify RGS3 and RGS2, as a transcriptional effector of this anti-lipolytic program. This finding provides a direct link between a STAT3-activating cytokine and a GPCR-driven kinase cascade. By promoting GTP hydrolysis on Gα proteins and inhibiting adenylyl cyclase, RGS proteins are positioned to dampen β-adrenergic signaling, cAMP generation and PKA-mediated HSL phosphorylation. The requirement for RGS3 and RGS3 in IL-10-mediated suppression of pHSL, together with STAT3-associated regulatory features near Rgs loci, supports a model in which IL-10 induces an RGS-dependent mechanism that limits catecholamine response without broadly affecting the lipolytic machinery.

These findings extend the role of adipose Tregs beyond their established function in suppressing inflammation. Adipose Tregs have been widely implicated in maintaining insulin sensitivity and tissue homeostasis, but their contribution has largely been interpreted through anti-inflammatory cytokine production and immune modulation. Our results suggest that Tregs also function as direct regulators of adipocyte signal transduction. In this model, Treg-derived IL-10 does not simply reduce inflammatory tone; it imposes a receptor-specific brake on adipocyte catecholamine sensitivity. This distinction is important because it positions immune cells as active regulators of lipid flux, rather than indirect modifiers of adipocyte function.

Together, these findings define a unique Treg–IL-10–IL10Rα–STAT3–RGS axis that restrains β-adrenergic activation of the PKA–HSL pathway (Fig. 4J). By identifying an inducible immune checkpoint for adipocyte lipolysis, this work reveals how adipose tissue integrates endocrine and immune signals to control lipid mobilization over time. More broadly, it suggests that immune cells do not merely buffer adipose inflammation, but directly shape the magnitude and duration of adipocyte intrinsic lipolytic pathways and lipid mobilization.

## Materials and Methods

### Cell Culture and Adipocyte Differentiation

Mouse C3H10T1/2 cells (CCL-226, ATCC) were maintained in Dulbecco’s Modified Eagle’s Medium (DMEM; MT10013CM, Corning) supplemented with 10% fetal bovine serum (FBS; FB-11, Omega Scientific) and 1% penicillin–streptomycin (MT30002CI, Corning) at 37 °C in a humidified incubator with 5% CO₂. Cells were routinely washed with phosphate-buffered saline (PBS; BP39920, Fisher) and detached using trypsin–EDTA (MT25053CI, Corning). Cells were passaged prior to reaching full confluence and were regularly tested to confirm the absence of mycoplasma contamination. For adipocyte differentiation, C3H10T1/2 cells were seeded at a density of 50,000 cells/mL in 6-well plates (07-200-83, Corning) with 2 mL of pre-adipogenic medium consisting of DMEM supplemented with 10% FBS, 1 nM triiodothyronine (T3; T2877, Sigma), and 5 µg/mL insulin (12-585-014, Gibco), and allowed to reach confluence. Differentiation was initiated (day 0) by replacing the growth medium with adipogenic induction medium containing DMEM supplemented with 10% FBS, T3, insulin, 2 µg/mL dexamethasone (D1756, Sigma), 0.5 mM 3-isobutyl-1-methylxanthine (IBMX; I7018, Sigma), 1 µM rosiglitazone (R2408, Sigma), and 125 µM indomethacin (I7378, Sigma), unless otherwise indicated. After 48 h, cells were switched to maintenance medium containing DMEM supplemented with 10% FBS, T3, insulin, and rosiglitazone. Maintenance medium was replaced every 2 days. Cells were harvested at the indicated time points for immunoblotting and RNA extraction.

### Human Participants and Cell Culture

Abdominal subcutaneous adipose tissue was obtained by needle aspiration from a healthy female volunteer (BMI: 33.7 kg/m², age: 30–40 years) with no history of diabetes, cancer, endocrine, or inflammatory diseases. Written informed consent was obtained prior to sample collection, and all procedures were approved by the Institutional Review Board of the Icahn School of Medicine at Mount Sinai. Isolated cells were cultured and allowed to differentiate into adipocytes. Cell lysates were collected 2 days post-confluence and after full differentiation (day 14 of culture).

During the maintenance phase, cells were refed every 2 days with freshly prepared maintenance medium consisting of DMEM:F12 supplemented with 10 nM insulin and 10 nM dexamethasone.

### Animals

C57BL/6 WT male and female mice (#000664) and Adipoq-Cre (#028020) mice were acquired from The Jackson Laboratory. TregIL10KO mice were generated using IL-10FL (#036598; B6.129P2(C)-Il10tm1Roer/MbogJ) and Foxp3YFP-cre (#016959; B6.129(Cg)-Foxp3tm4(YFP/icre)Ayr/J) mice. iAdIL10RαKO mice were generated using IL10Raflox (#028146; B6(SJL)-Il10ratm1.1Tlg/J), tetO-cre (#006234; B6.Cg-Tg(tetO-cre)1Jaw/J and Adipoq-rtTA (#033448; C57BL/6-Tg(Adipoq-rtTA)2Zvw/J) mice. All mice were maintained in a pathogen-free, barrier-protected environment (12:12 h light/dark cycle, 22–24°C) at the Mount Sinai animal facilities. Mice were fed either a chow diet (Purina 5001), a 60% high-fat diet (HFD; Research Diets), or a doxycycline chow diet containing 625 mg/kg doxycycline (TD.01306; Inotiv), with ad libitum access to food and water. Animal experiments were conducted in accordance with the guidelines of the Mount Sinai Institutional Animal Care and Use Committee. Randomization was not used in the allocation process because experimental groups were primarily separated based on genotype or sex.

### Pharmacological Stimulation

Where indicated, differentiated adipocytes were stimulated with 5 µM CL-316,243 (C5976, Sigma) or vehicle control for 15 min, or with 100 ng/mL interleukin-10 (IL-10) (Sino Biological) for 3 h. For combined treatments, cells were first treated with IL-10 (100 ng/mL) for 3 h, followed by stimulation with CL-316,243 (5 µM) for 15 min.

### Single-cell RNA-sequencing (scRNA-seq)

Adult male C57BL/6J mice (10–12 weeks) were administered intraperitoneal injections of the β3-adrenergic agonist CL-316,243 (1 mg kg^-1^ in sterile saline) and sacrificed at 0 h (D0, vehicle baseline), 24 h (D1) or 48 h (D2) post-injection. Inguinal white adipose tissue (iWAT) depots were excised, minced and digested in HBSS containing 1 mg ml^-1^ collagenase type II (Worthington) and 2% BSA for 30 min at 37 °C with gentle shaking. Digests were quenched with cold DMEM/10% FBS, passed through a 100-µm strainer, and floating mature adipocytes were removed by centrifugation at 500 × *g* for 5 min. The stromal-vascular fraction (SVF) pellet was treated with ACK lysis buffer for 3 min, washed, and resuspended in PBS / 0.04% BSA. Viability (≥85%) and cell concentration were confirmed on a Countess II using trypan blue.

### 10x Genomics library preparation and sequencing

Approximately 10,000 viable SVF cells per sample were loaded onto a Chromium Controller using the Chromium Next GEM Single Cell 3’ Reagent Kit v3.1 (10x Genomics) according to the manufacturer’s protocol. Reverse transcription, cDNA amplification (12 cycles) and library construction were performed as recommended. Libraries were quantified by Qubit and Bioanalyzer, pooled, and sequenced on an Illumina NovaSeq 6000 (paired-end) to a target depth of ≥30,000 reads per cell.

### Read alignment and count-matrix generation

Raw BCL files were demultiplexed with ‘cellranger mkfastq’ and aligned to the mouse reference genome (refdata-gex-mm10-2020-A; GENCODE vM23/Ensembl 98) with ‘cellranger count’ (10x Genomics, v7.1.0). Per-sample filtered feature–barcode matrices were retained for downstream analysis.

### Quality control, doublet detection and filtering

Filtered matrices were loaded into Python (scanpy v1.10) as a unified AnnData object spanning the three time points (D0, D1, D2). Per-cell quality metrics were computed with ‘sc.pp.calculate_qc_metrics’. Cells were retained that expressed ≥200 unique genes (‘n_genes_by_counts’ ≥ 200) and had mitochondrial transcript fraction < 15% (‘pct_counts_mt’ < 15). Putative doublets were flagged per sample with Scrublet (default expected_doublet_rate = 0.06) and removed. Median post-QC metrics were 1,650 / 1,938 / 1,681 detected genes and 4.1% / 3.9% / 8.4% mitochondrial reads at D0 / D1 / D2, yielding a final matrix of \**22,826 cells × 33,696 genes*\* (D0 = 5,942; D1 = 7,645; D2 = 9,239).

### Normalization, feature selection and dimensionality reduction

Raw counts were preserved in ‘adata.layers[’counts’]’. Counts were library-size–normalized to 10,000 per cell (‘sc.pp.normalize_total’, target_sum = 1e4) and natural-log transformed with a pseudo-count of 1 (‘sc.pp.log1p’). The top 3,000 highly variable genes were selected per time point with the seurat_v3 (‘sc.pp.highly_variable_genes(flavor=’seurat_v3’, n_top_genes=3000, batch_key=’sample_id’)’) and used for downstream embedding. Expression values were scaled (zero mean, unit variance; clip = 10) and a 50-component PCA was computed on the scaled HVG matrix with ‘sc.tl.pca(svd_solver=’arpack’, random_state=42)’.

### Batch correction with Harmony

To remove run-level technical variance across the three captures while preserving the biological CL response, Harmony integration (‘scanpy.external.pp.harmony_integratè, harmonypy v0.0.9) was run on the first 30 PCs with ‘sample_id’ as the batch key (‘theta = 2’, ‘max_iter_harmony = 20’, ‘random_state = 42’). Harmony embeddings were written to ‘adata.obsm[’X_pca_harmony’]’; the pre-correction PCA was retained as ‘adata.obsm[’X_pca’]’ and a pre-Harmony UMAP was stored as ‘adata.obsm[’X_umap_pre_harmony’]’ for diagnostic comparison.

### Neighbour graph, UMAP and Leiden clustering

A k-nearest-neighbour graph was built on the Harmony embedding with ‘sc.pp.neighbors(use_rep=’X_pca_harmony’, n_neighbors=30, n_pcs=30, metric=’euclidean’, random_state=42)’. UMAP coordinates were computed with ‘sc.tl.umap(min_dist=0.3, spread=1.0, random_state=42)’. Communities were called with the Leiden algorithm (‘sc.tl.leiden’, leidenalg v0.10) at two resolutions — 0.5 (15 clusters, used as the primary partition) and 1.0 (21 clusters, used for fine sub-population resolution). Cluster identities were stable across 10 seeds (ARI > 0.92).

### Cell-type annotation by marker scoring and reference label transfer

Cluster identities were assigned by an orthogonal two-step procedure. (i) \**Marker-score annotation: curated lineage marker panels (B cell:* Cd19, Ms4a1, Cd79a*; CD4 T:* Cd3e, Cd4*; CD8 T:* Cd3e, Cd8a, Cd8b1*; NK:* Klrb1c, Ncr1, Nkg7*; macrophage:* Adgre1, Cd68, Lyz2*; dendritic:* Itgax, Flt3*; neutrophil:* S100a8, S100a9*; endothelial:* Pecam1, Cdh5, Kdr*; stromal/APC:* Pdgfra, Dpp4, Ly6a*; T_reg:* Foxp3, Il2ra, Ctla4) were converted to z-scored module scores with ‘sc.tl.score_genes’. (ii) Reference-based label transfer: cluster centroids were matched to the Loft et al. 2025 ^28^ consensus mouse-adipose atlas using a logistic-regression classifier trained on the reference (scaled log-normalised, balanced classes, L2-regularised, 5-fold CV; confidence stored in ‘adata.obs[’conf_score’]’). Final labels (‘cell_type_final’, 13 categories) required concordance between the two methods and clusters that disagreed were marked ‘unknown’ (1,865 cells, 8.2%).

### Treg and IL-10⁺ Treg gating

Within the lymphoid compartment, regulatory T cells were defined *in silico* as ‘is_treg = Truè when cells co-expressed *Cd3e* > 0 and *Foxp3* > 0 (log-normalised counts) and were assigned ‘cell_type_final == ’Treg’’. Treg-like cells expressing *Foxp3* outside the CD3^+^ lymphoid lineage were retained in a separate category (‘Foxp3_nonT’, n = 26) to avoid contamination of the lineage-restricted Treg pool. IL-10–producing Tregs (‘is_il10_treg’) were defined as Tregs (above) with *Il10* > 0. The final pools were 4,446 CD3^+^ T cells, 67 lineage-restricted Tregs and IL-10^+^ Tregs across the three time points.

### Differential abundance across the time course

Per–cell-type proportions were computed at D0, D1 and D2 and compared using a generalised linear model with a quasi-binomial link on cell counts (offset = log[total cells per sample]) implemented via *statsmodels*. Two-sided *P* values were Benjamini–Hochberg-corrected across all cell types; differential abundance was reported as log₂ fold-change vs D0 with 95% confidence intervals.

### STAT3 and JAK Inhibitor Treatment in 10T1/2 Cells and Human adipocytes

10T1/2 cells or human ADSCs were cultured and differentiated into adipocytes as described above. To assess the role of JAK–STAT signaling, cells were treated with STAT3 Inhibitor III, WP1066 (MilliporeSigma/Calbiochem; CAS 857064-38-1) or InSolution™ JAK Inhibitor I (Calbiochem; CAS 457081-03-7). Inhibitors were prepared according to the manufacturers’ instructions and diluted into culture medium at the indicated working concentrations. Control cells were treated with an equivalent volume of vehicle. Cells were treated for the indicated time points, after which they were harvested for downstream analyses, including protein extraction for immunoblotting. Cell morphology was monitored during treatment to assess gross cytotoxicity.

### Extracellular Flux Analysis

Oxygen consumption rate (OCR) was measured using a Seahorse XFe96 Extracellular Flux Analyzer (Agilent) according to the manufacturer’s instructions. 10T1/2 cells were seeded in Seahorse XFe96 cell culture microplates (Agilent) at a density of 3,000 cells per well in 150 µL complete DMEM supplemented with 10% FBS and differentiated into adipocytes prior to analysis. Before the assay, fully differentiated adipocytes were incubated in glucose-free medium for 3 hours, followed by treatment with interleukin-10 (IL-10; 100 ng/mL) for 3 hours. Control cells were maintained in standard complete medium. Where indicated, cells were treated with CL-316,243 (CL; 5 µM) prior to Seahorse analysis. On the day of the assay, cells were washed three times and incubated in 180 µL XF Base Medium (Agilent) supplemented with 25 mM glucose, 2 mM sodium pyruvate, and 2 mM glutamine, adjusted to pH 7.4. Cells were equilibrated for 45 minutes at 37°C in a non-CO₂ incubator before measurement. For the mitochondrial stress test, three baseline OCR measurements were obtained, followed by sequential injections of oligomycin (2 µM), FCCP (2 µM), and rotenone/antimycin A (0.5 µM). OCR parameters were calculated using the Seahorse XF Cell Mito Stress Test Report Generator (Agilent) and normalized to total protein.

### Phosphoproteomic Analysis

Phosphoproteomic analysis was performed as a service by Creative Proteomics using cell pellet samples. Briefly, six cell pellet samples were lysed in buffer containing 8 M urea supplemented with protease and phosphatase inhibitors. Samples were sonicated and centrifuged at 12,000 × g for 10 minutes at 4°C, and protein concentration was determined using a BCA assay. Proteins were digested using a filter-aided sample preparation approach. Protein samples were reduced with 10 mM DTT at 56°C for 1 hour and alkylated with 20 mM iodoacetamide at room temperature in the dark for 1 hour. Proteins were then digested overnight at 37°C with trypsin at an enzyme-to-protein ratio of 1:50. Phosphorylated peptides were enriched using Fe-IMAC beads. Enriched phosphopeptides were eluted, acidified with trifluoroacetic acid, dried using a SpeedVac, and resuspended in 0.1% formic acid prior to analysis. Samples were analyzed by nano-LC-MS/MS using an Ultimate 3000 nano UHPLC system coupled to a Q Exactive HF mass spectrometer with an electrospray ionization nanospray source. Peptides were separated on C18 trapping and analytical columns using a linear acetonitrile gradient at a flow rate of 250 nL/min. MS data were acquired in data-dependent mode, with full scans collected over 300–1,650 m/z and MS/MS analysis performed using a Top 20 method. Raw mass spectrometry files were analyzed using MaxQuant. Searches were performed against the appropriate protein database with trypsin specificity and up to two missed cleavages allowed. Carbamidomethylation of cysteine was set as a fixed modification, while methionine oxidation and phosphorylation of serine, threonine, and tyrosine were set as variable modifications. Differentially regulated phosphoproteins/phosphorylation sites were defined using a fold-change threshold of >1.5 for upregulation and <0.67 for downregulation.

### RNA-seq Analysis

Total RNA was extracted from cultured cell samples as described previously. Strand-specific RNA-seq libraries were prepared from 500 ng of total RNA using the TruSeq Stranded Total RNA Library Prep Kit according to the manufacturer’s protocol. Libraries were sequenced as 50-bp single-end reads on an Illumina HiSeq 2000 or HiSeq 4000 instrument. Reads were mapped to the mouse reference genome NCBI37/mm9 with TopHat v1.3.3, permitting a single alignment per read and allowing up to two mismatches. Gene-level expression was calculated as RPKM/FPKM using the mRNA quantification pipeline in SeqMonk. For in vitro experiments, expression values were averaged from three independent biological replicates used for library generation. Genes were retained for downstream analyses if they had a maximum FPKM of at least 10 at any time point and were longer than 200 bp. Differential expression was assessed in R/Bioconductor using DESeq2, and multiple-testing correction was performed using the Benjamini–Hochberg method.

### ATAC-seq Library Preparation

ATAC-seq libraries were prepared from 10T1/2 cells using the Zymo-Seq ATAC Library Kit (Zymo Research) according to the manufacturer’s protocol. Briefly, 10T1/2 cells were harvested, counted, and processed using the recommended cell input for the kit. Cells were gently lysed to isolate nuclei, followed by tagmentation of accessible chromatin regions using the supplied transposase reaction reagents. Following tagmentation, DNA fragments were purified and amplified by index PCR using the kit-provided library amplification reagents and unique dual-index primers. Amplified libraries were purified according to the manufacturer’s instructions and assessed for concentration and quality prior to sequencing. Library quality was evaluated based on DNA concentration and fragment size distribution. Prepared ATAC-seq libraries were submitted for next-generation sequencing. The Zymo-Seq ATAC workflow includes gentle cell lysis and nuclei preparation, tagmentation of open chromatin regions, and indexed PCR amplification to generate sequencing-ready libraries from mammalian cells. The kit is designed to prepare ATAC-seq libraries from low-input samples, with recommended input as low as 50,000 cells.

### Body Composition Measurements

Body composition, including fat mass and lean mass, was measured using an EchoMRI Body Composition Analyzer.

### RGS2 and RGS3 siRNA Transfection in 10T1/2 Cells

10T1/2 cells were cultured under standard conditions and transfected with mouse RGS2 and RGS3 siRNA (Santa Cruz Biotechnology) to knock downRGS2 or RGS3 expression. The RGS2 or RGS3 siRNA reagent consists of a pool of target-specific siRNAs designed against mouse RGS2 or RGS3. Transfections were performed using Lipofectamine transfection reagent (Thermo Fisher Scientific) according to the manufacturer’s instructions. Briefly, cells were seeded to reach the appropriate confluency at the time of transfection. RGS2 and RGS3 siRNA or non-targeting control siRNA was diluted in serum-free medium and mixed with Lipofectamine reagent. After incubation to allow complex formation, the siRNA–Lipofectamine mixture was added to cells. Following transfection, cells were maintained under standard culture conditions for the indicated time period before downstream analysis. Knockdown efficiency was assessed by measuring RGS2 and RGS3 protein levels. Cells were then harvested for subsequent analyses, including RNA extraction, protein lysate preparation, and functional assays as indicated. Non-targeting siRNA-transfected cells were used as controls.

### RGS3 shRNA Lentiviral Transduction

10T1/2 cells were transduced with mouse RGS3 shRNA lentiviral particles (RGS3 shRNA [m] Lentiviral Particles, Santa Cruz Biotechnology, sc-40662-V) to achieve stable knockdown of RGS3 expression. Cells were plated 24 hours prior to transduction to reach approximately 50% confluency at the time of infection. On the day of transduction, culture medium was replaced with complete medium containing Polybrene (Thermo Fisher Scientific) at a final concentration of 4 µg/mL**, a**nd RGS3 shRNA lentiviral particles were added directly to the cells. Cells were gently mixed and incubated overnight under standard culture conditions. The following day, viral-containing medium was removed and replaced with fresh complete medium without Polybrene. For stable knockdown, cells were expanded 24–48 hours post-transduction and selected with puromycin (Thermo Fisher Scientific) at a final concentration of 4 µg/mL. Puromycin-containing medium was replaced every 3–4 days until resistant cell populations were established. Knockdown efficiency was confirmed by assessing RGS3 mRNA protein levels. Non-targeting control shRNA lentiviral particles were used as controls.

### Generation of *Rgs3* knockout 10T1/2 cells by CRISPR–Cas9

*Rgs3* knockout C3H/10T1/2 cells were generated via CRISPR–Cas9 ribonucleoprotein (RNP) electroporation. Cells were maintained in Dulbecco’s Modified Eagle Medium (DMEM) containing 10% fetal bovine serum (FBS) and 1% penicillin–streptomycin (Thermo Fisher Scientific) at 37°C with 5% CO₂. Two single guide RNAs (sgRNAs) targeting *Rgs3* were designed in silico and synthesized by GenScript (sgRNA#1: 5′-ACAACUGAAUGAGAGACCCG-3′; sgRNA#2: 5′-ACGAGCAGGAUGAUCUCGCU-3′). Prior to RNP complex formation, the sgRNAs were pooled to enhance knockout efficiency. Recombinant Cas9 protein (TrueCut Cas9 Protein v2, Cat# A36498; Invitrogen) was then complexed with the pooled sgRNAs at a 1.2:1 molar ratio in Resuspension Buffer R (Thermo Fisher Scientific) for 15 min at room temperature. Cells were detached with trypsin–EDTA (Invivogen), washed with DPBS (Gibco), and resuspended in Neon Resuspension Buffer R. Ten microliters of the cell suspension was combined with RNP complexes and electroporated using the Neon Transfection System (Thermo Fisher Scientific) under the following conditions: 1,300 V, 20 ms, 2 pulses. Immediately after electroporation, cells were transferred into pre-warmed complete medium (DMEM with 10% FBS) and maintained under standard conditions, with medium replacement at 12–24 h. Once sufficiently expanded, genomic DNA was extracted by a standard lysis-based method; the target locus was amplified by PCR and submitted for Sanger sequencing (Genewiz) to verify indel formation and *Rgs3* knockout status. For clonal isolation, cells were diluted and seeded at approximately one cell per well in 96-well plates (Corning), and individual clones were expanded for genomic DNA extraction and genotyping by PCR–Sanger sequencing to confirm *Rgs3* knockout.

### Regulatory T Cell analysis and flow cytometry

For flow cytometry analysis, blood was collected after a 6-hour fast by cardiac puncture using a 23G needle and a 1 mL syringe pre-flushed with 0.5 M EDTA. Following blood collection, mice were euthanized with isoflurane and perfused with 1x phosphate-buffered saline (PBS). Both inguinal (iWAT) and epididymal white adipose tissue (eWAT) depots were harvested and rinsed in cold phosphate-buffered saline; inguinal lymph nodes were removed via ex vivo dissection. Whole adipose tissue depots (∼100-300 mg) were then minced in 1 mL of digestion buffer (450 U/mL collagenase I, 125 U/mL collagenase XI, 60 U/mL DNase I, and 60 U/mL hyaluronidase in 1x PBS) and incubated at 37°C in a shaking incubator at 1000 rpm for 40 minutes. Once digested, tissues were filtered through a 70 µm cell strainer into 50-mL conical tubes and washed once in PBS; single-cell suspensions were then pelleted by centrifugation at 1800 RPM for 3-5 minutes at 4°C. Pellets from blood and adipose tissue depots were resuspended in 1mL of RBC lysis buffer (BioLegend #420301) to lyse erythrocytes and incubated at room temperature for 10 minutes, and quenched with 9 mL of PBS. This process was repeated for blood.

Single-cell suspensions were centrifuged for 3-5 minutes at 1800 RPM at 4°C, then resuspended in an extracellular staining mix prepared in fresh MACS buffer (0.5% bovine serum albumin and 2 mM EDTA in PBS) for 30-45 minutes at 4°C. To differentiate between live and dead cells, adipose and blood cell suspensions were stained with 200 microliters of Live/Dead Blue (Thermo Fisher #L23105) prepared in 1x PBS at a 1:1000 dilution, for 20 minutes at 4°C, and quenched with PBS. The following antibodies were used for extracellular flow cytometry analysis at a dilution of 1:700: CD45-BV711 (BioLegend #103147), CD11b-APC-Cy7 (BioLegend #101225), CD90.2-Alexa Fluor 700 (BioLegend #105319), CD3-PE (BioLegend #100308), B220-BUV805 (Invitrogen #368-0452-80), CD4-BV785 (BioLegend #100552), CD8b- BUV737 (BD Bioscience #741811), CD25-BV421 (BioLegend #102043). Samples were then quenched with 1X PBS, fixed and permeabilized using Foxp3 fixation/permeabilization working solution (Invitrogen #00-5523-00) for 45 minutes at 4°C, protected from light, and quenched with 1X Permeabilization Buffer, per the manufacturer’s instructions. Pellets were resuspended in an intracellular/intranuclear staining mix prepared in 1x Permeabilization buffer, incubated at room temperature protected from light for 45 minutes, washed twice in 2mL of 1x Permeabilization Buffer, and finally resuspended in 300 microliters of fresh MACS buffer, then stored at 4°C protected from light. The following antibodies were used for intranuclear/intracellular flow cytometry analysis at a dilution of 1:200: FOXP3-Alexa Fluor 647 (BioLegend #320014), IL10-FITC (BioLegend #505005). Samples were acquired using the Cytek Aurora with Spectroflow (Cytek) and analyzed with FlowJo software (BD Biosciences). Cell populations were quantified using precision counting beads (BioLegend #424902) and the gating strategy was set up using Fluorescence Minus One (FMO) controls. Cells were identified according to surface and intra-nuclear/cellular markers listed in Supplementary Table 2.

### CL-316,243 Treatment in Mice

Male wild-type mice aged 8–10 weeks were used for all experiments. Mice were randomly assigned to treatment groups receiving either CL-316,243 (CL) or phosphate-buffered saline (PBS) as control. CL-316,243 was administered via intraperitoneal injection at the indicated conditions. For acute stimulation experiments, mice received a single injection of CL and were sacrificed 30 minutes post-injection. For chronic stimulation, mice were treated with CL for 48 hours, receiving injections every 24 hours during this period. Control animals received PBS injections following the same schedule. At the indicated time points, mice were euthanized, and tissues were collected for downstream analyses.

### Doxycycline Diet for iAdIL10RαKO mice

A doxycycline chow diet containing 625 mg/kg diet (TD.01306; Inotiv) was used. This diet was designed to deliver a daily dose of 2–3 mg of doxycycline based on an average mouse consumption of 4–5 g/day. Doxycycline hyclate contains approximately 87% doxycycline. Mice were fed the doxycycline diet for 7 days, as this provides a stable and non-stressful method for inducing gene expression in animal models. After 7 days, the diet was replaced with a normal chow diet for the next 21 days. Organs were then harvested following PBS and CL injections administered 30 minutes and 48 hours prior to tissue collection.

### Glucose Tolerance Test (GTT) and Insulin Tolerance Test (ITT)

Glucose tolerance tests (GTT) and insulin tolerance tests (ITT) were performed to assess systemic glucose homeostasis and insulin sensitivity in mice. For both assays, mice were fasted for 6 hours prior to testing. For the GTT, mice received an intraperitoneal injection of glucose (D-(+) Glucose Solution, G8769-100ml, Sigma Aldrich, Lot# RNBK9807) at a dose of 2 g/kg body weight. Blood glucose levels were measured at the indicated time points using an ACCU-CHEK Active glucometer (Roche). For the ITT, following the same fasting period, mice were administered insulin (Insulin Human injection 100units/ml (U-100), Novolin R NDC 0169-1833-02) via intraperitoneal injection at a dose of 1 U/kg body weight. Blood glucose levels were measured at the indicated time points using the same glucometer. All measurements were performed according to standard procedures, and glucose values were recorded for subsequent analysis.

### Immunoblotting

Cells or tissues were lysed in ice-cold RIPA buffer (Boston Bioproducts, NC9923398) supplemented with protease and phosphatase inhibitors (Thermo Fisher Scientific, PIA32961PM). Lysates were incubated on ice for 5 minutes and clarified by centrifugation at 13,000 × g for 15 minutes at 4°C. Protein concentration was determined using a BCA assay (Thermo Fisher Scientific, PI23227). Equal amounts of protein were mixed with LDS sample buffer (Invitrogen, B0007) and 50 mM dithiothreitol (Thermo Fisher Scientific, FERR0861), followed by denaturation at 100°C for 15 minutes. Proteins were separated by SDS–PAGE and transferred onto PVDF membranes (Sigma, IPFL85R). Membranes were blocked using blocking buffer (LI-COR, 927-60010) and incubated with primary antibodies overnight at 4°C. After washing, membranes were incubated with fluorescent secondary antibodies for 1 hour at room temperature. Protein signals were detected using a LI-COR Odyssey imaging system and quantified using Empiria Studio software. Protein expression levels were normalized to housekeeping proteins, including tubulin and GAPDH, as indicated in the figure legends. All antibodies used in this study are listed in Supplementary Table 1.

### Quantification and Statistical Analysis

Data are presented as mean ± S.E.M. unless otherwise indicated. Experiments were performed with independent biological replicates, as specified in the figure legends. Sample sizes were determined based on prior experimental experience, and no statistical methods were used to predetermine sample size. Samples were excluded only based on predefined technical criteria, including failed infection, poor RNA or protein quality, or instrument malfunction. Statistical analyses were conducted using GraphPad Prism. Comparisons between two groups were performed using two-tailed unpaired Student’s t-tests, or paired tests when appropriate. For multiple group comparisons, one-way or two-way ANOVA with appropriate post hoc multiple comparison correction was used. Repeated-measures analyses were applied for longitudinal data where applicable. A p-value < 0.05 was considered statistically significant.

## Acknowledgements

Lipidomics was performed by Dr. Kevin Williams at the UCLA Lipidomics Core. Phosphoproteomics was performed by Creative Proteomics. hADSCs were gifts from Dr. Susan Fried (ISMMS). P.R is supported by R01DK136035, DP1DK140003, Irma T. Hirschl/Monique Weill-Caulier Foundation Scholar Award. L.G. is supported by AHA 25TPA1480256. C.H.C supported by F32DK141191. N.F.B supported by T32HL176457. The funders had no role in study design, data collection and interpretation, or the decision to submit the work for publication.

## Competing interests

The authors declare no competing interests

Supplementary Figures

## Supplementary Figures

**Fig. S1.**
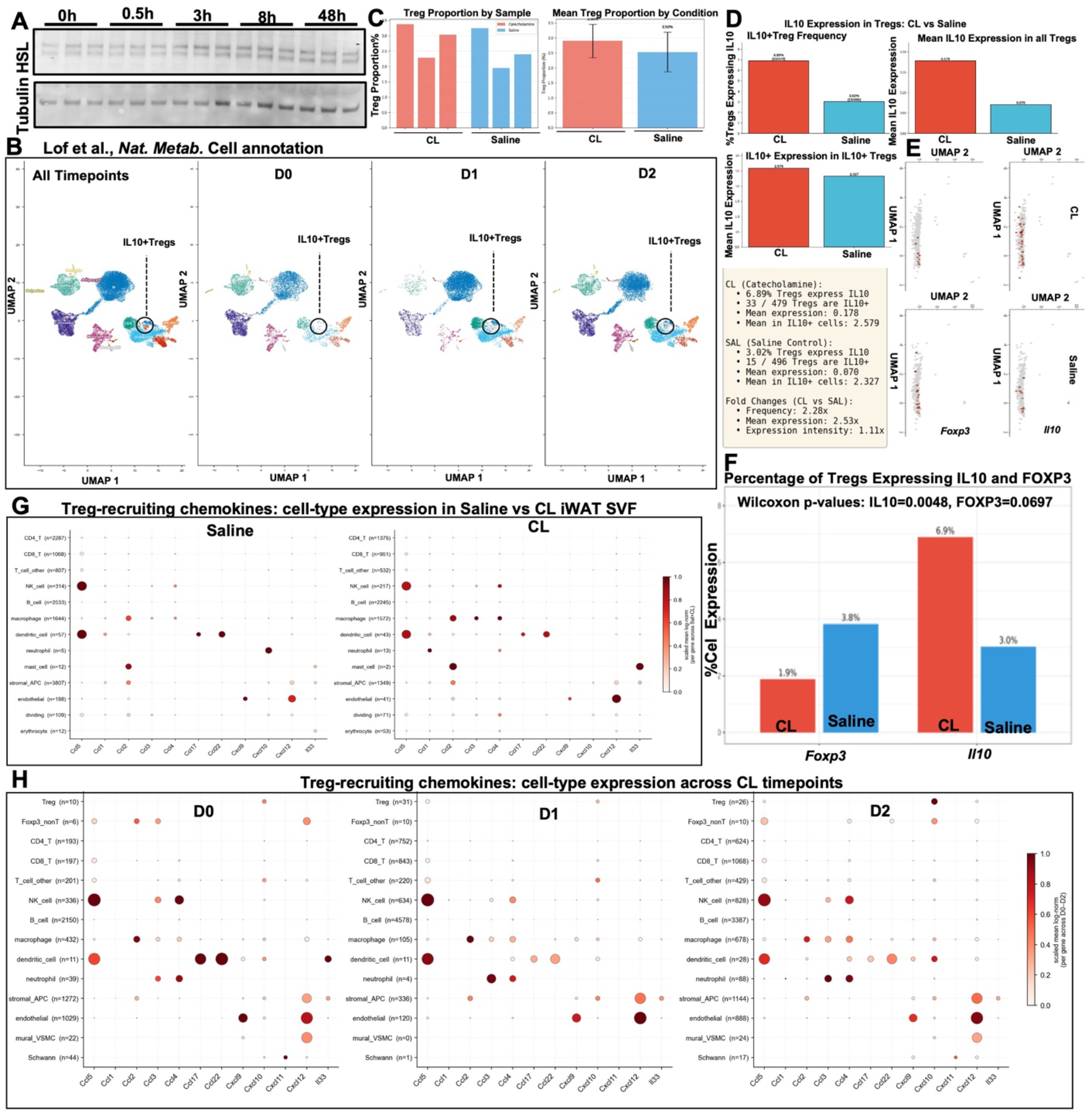
Time-course of CL-induced IL-10 expression and Treg abundance. (A) HSL and Tubulin western blots across the CL time course. (B) UMAP of all iWAT SVF cells colored by LofT et al. cell annotation across all time points and D0, D1, D2; IL-10+ Tregs circled. (C) Per-sample and mean Treg proportion in CL versus saline. (D) IL-10 expression metrics in Tregs (IL10+ Treg frequency, mean IL-10 expression in all Tregs, mean IL-10 expression in IL10+ Tregs) with summary statistics (CL: 6.89% IL10+, mean 0.178; SAL: 3.02% IL10^+, mean 0.070; frequency FC 2.28×, mean-expression FC 2.53×). (E) Foxp3 and Il10 UMAP feature plots in CL and saline. (F) Percentage of Tregs expressing IL10 and FOXP3 with Wilcoxon p-values (IL10 P = 0.0048, FOXP3 P = 0.0697). (G) Dot plots of Treg-recruiting chemokines (Ccl1, Ccl17, Ccl22, Cxcl12 and others) across cell types in saline and CL iWAT SVF (color, scaled mean expression; size, fraction of cells expressing). (H) The same dot plots resolved by CL time point (D0, D1, D2). n cell counts per cell type indicated.

**Fig. S2.**
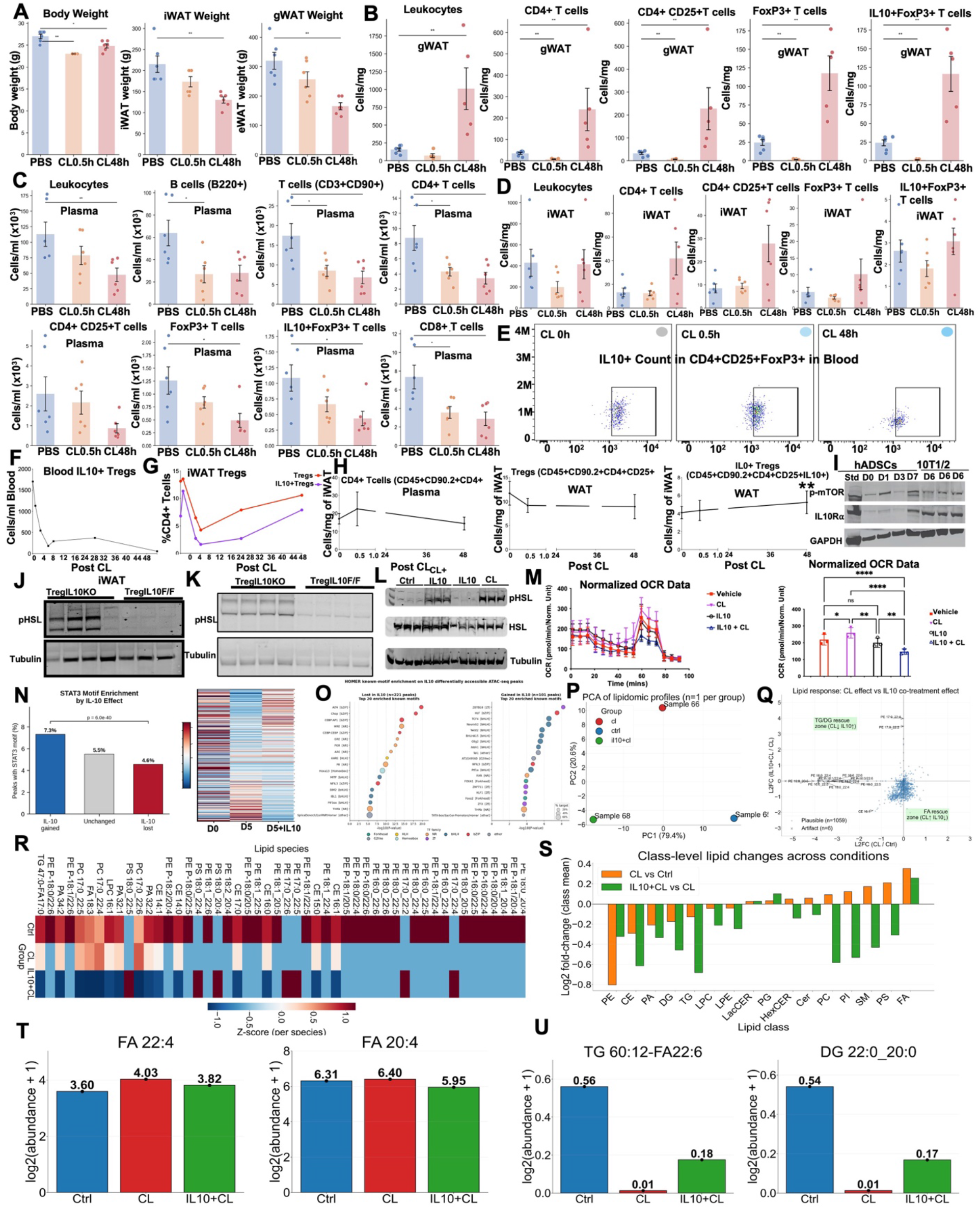
IL-10 from Tregs restrains adrenergic activation of HSL. (A) Body weight, inguinal WAT (iWAT) weight, and epididymal WAT (eWAT/gWAT) weight in C57BL/6J mice 0.5 h or 48 h after injection of CL-316,243 versus PBS (n = 5–6 per group). Mean ± s.e.m.; P < 0.05, P < 0.01 by one-way ANOVA. (B) Absolute immune-cell densities (cells per mg of tissue) in gWAT for total leukocytes, CD4+ T cells, CD4+CD25+ T cells, Foxp3+Tregs, and IL-10+Foxp3+Tregs across PBS, CL 0.5 h, and CL 48 h. P < 0.01 by one-way ANOVA. (C) Absolute densities (cells per ml × 10^3) of plasma leukocytes, B220^+^ B cells, CD3+CD90+ T cells, and CD4+T cells across the same time course. P < 0.05, *P < 0.01. (D) Absolute densities (cells per mg) of leukocytes, CD4+ T cells, CD4+CD25+, Foxp3+ Tregs and IL-10+Foxp3+ Tregs in iWAT. (E) Representative FACS dot plots (FSC-H × FITC–IL-10) of the IL-10+ gate within blood CD4+CD25+Foxp3+Tregs at CL 0 h, 0.5 h and 48 h. (F) Plasma densities (cells per ml × 10^3) of IL-10+Tregs across the time course. (G) Densely sampled time course (0–48 h post-CL) of %CD4+ T cells WAT Tregs or IL-10+ Tregs. (H) Compartment-matched time courses (0–48 h) of plasma CD4+ T cells, WAT CD4+CD25+ Tregs (CD45+CD90.2+CD4+CD25+) and WAT IL- 10+ Tregs (CD45+CD90.2+CD4+CD25+IL-10+); **P < 0.01. (I) Western blot of phospho-mTOR, IL-10Rα and GAPDH across hADSC and 10T1/2 differentiation time points (D0, D1, D3, D7) with a D6/D6/D6 endpoint triplicate. (J-L) Western blot of pHSL, total HSL and tubulin in primary adipocytes treated with Ctrl, CL+IL-10, IL-10 or CL (L); parallel pHSL blots in iWAT (J) and gWAT (K) lysates from TregIL10^KO^ versus TregIL10^F/F^ mice. (M) Normalized OCR (right): real-time oxygen consumption rate (pmol/min per normalized unit) across Vehicle, CL, IL-10 and IL-10+CL with summary bar graph; n = 3 per group; P < 0.05, *P < 0.01, P < 0.0001 by one-way ANOVA. (N) Bar plot of the percentage of ATAC-seq peaks containing a STAT3 motif partitioned by IL-10 effect — IL-10 gained (7.3%), Unchanged (5.5%), IL-10 lost (4.6%); P* = 6.0 × 10−40, two-sided Fisher’s exact test. Accompanying heatmap displays STAT3- footprinted peak signal at D0, D5 and D5+IL-10. (O) HOMER known-motif enrichment on IL-10–differentially accessible ATAC-seq peaks: top 20 motifs lost in IL-10 (n = 221 peaks, left) and gained in IL-10 (n = 101 peaks, right); bubble size, % target sequences; color, TF family. (P) Principal-component analysis (PCA) of untargeted lipidomic profiles (n = 3 combined and 1 replicate per group; Sample 65 = ctrl, 66 = CL, 68 = IL-10+CL); axes show PC1 (79.4%) and PC2 (20.6%). (Q) Per-lipid scatter plot of log2FC IL-10+CL/CL (y-axis) versus log2FC CL/Ctrl (x-axis); shaded green regions define the TG/DG rescue zone (CL↓ / IL-10↑, upper-left) and FA rescue zone (CL↑ / IL-10↓, lower-right). Plausible species, n = 1,059; artifacts excluded, n = 6. (R) Per-species z-score heatmap of selected differentially abundant lipids across Ctrl, CL and IL-10+CL. (S) Class-level log2 fold-change (class-mean) for the CL/Ctrl (orange) and IL-10+CL/CL (green) comparisons across 16 lipid classes (PE, CE, PA, DG, TG, LPC, LPE, LacCER, PG, HexCER, Cer, PC, PI, SM, PS, FA). (T) Absolute abundances [log2(abundance + 1)] of FA 22:4 and FA 20:4 across Ctrl, CL and IL-10+CL. (U) Absolute abundances [log2(abundance + 1)] of TG 60:12-FA22:6 and DG 22:0_20:0 across Ctrl, CL and IL-10+CL. All bar plots show mean ± s.e.m.; sample sizes per group are indicated in panels.

**Fig. S3.**
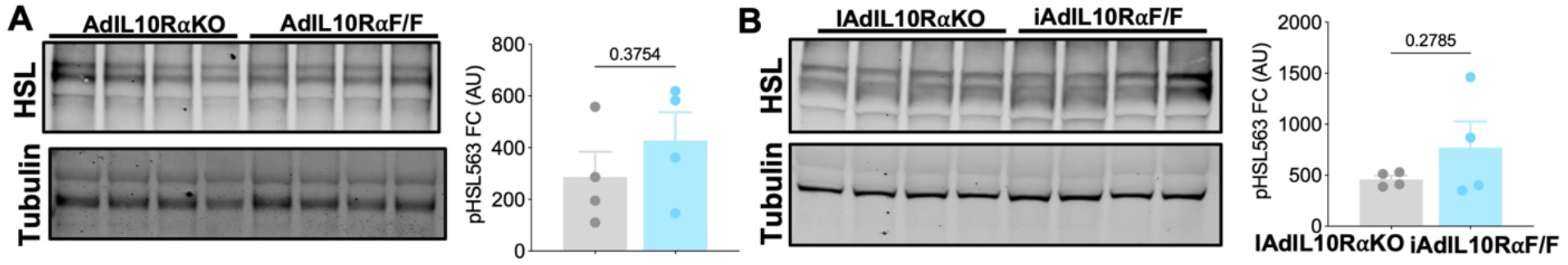
IL-10Rα in adipocytes is required to mediate the anti-lipolytic effect of Treg-IL10. (A and B) Total HSL and Tubulin western blots and HSL quantification in WATs of iAdIL10RKO versus iAdIL10RF/F and iAdIL10RKO versus iAdIL10RF/F.

**Fig. S4.**
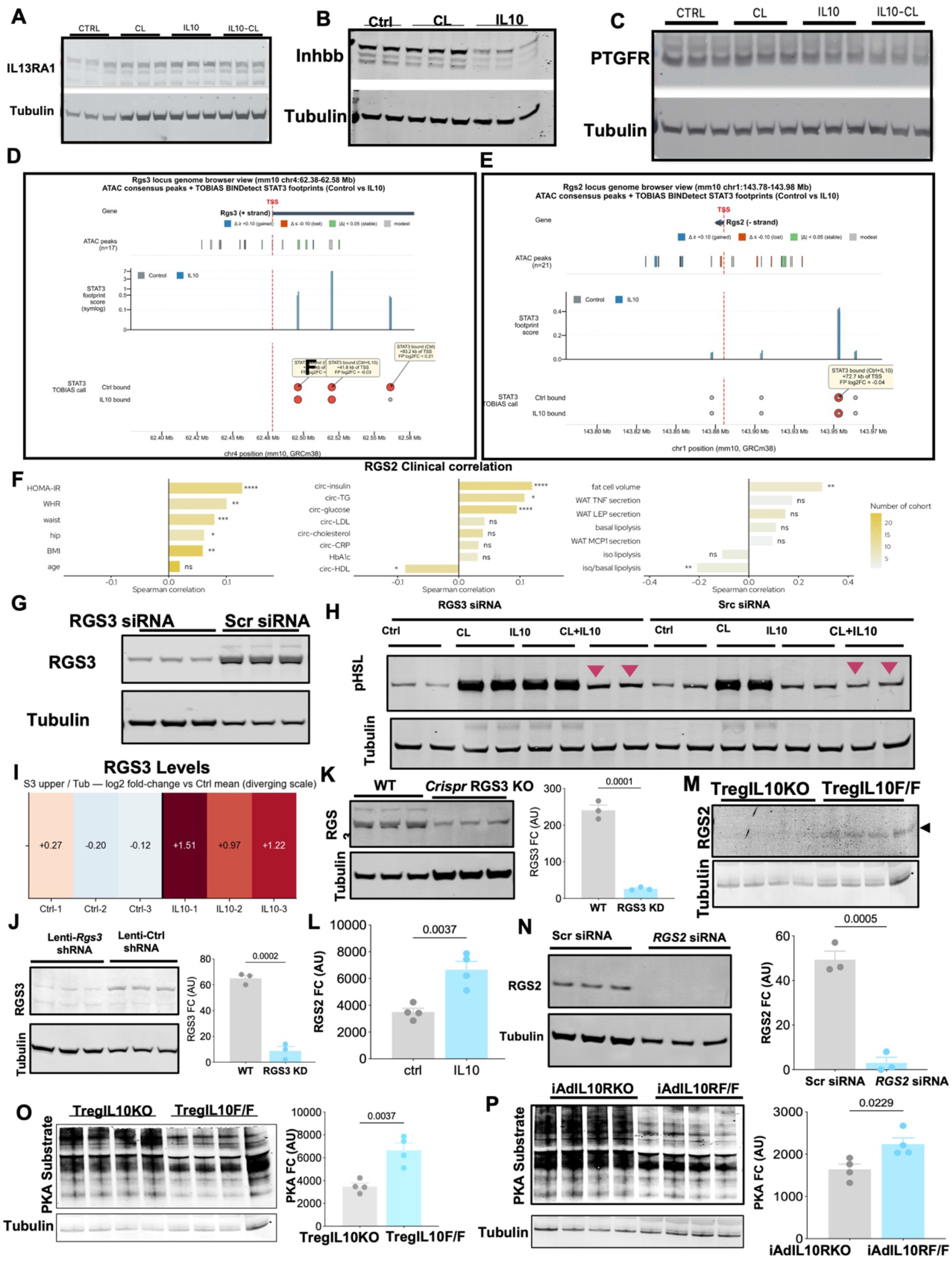
IL10 activated RGS3 and RGS2 control HSL activation. (A–C) IL13RA1, INHBB and PTGFR western blots in differentiated adipocytes across Ctrl, CL, IL-10 and CL+IL-10 conditions. (D and E) Rgs3 and Rgs2 locus genome-browser views (mm10) showing ATAC consensus peaks, gain/loss/stable peak status, STAT3 footprint scores and TOBIAS BINDetect STAT3-bound calls in Control versus IL-10; orange labels mark IL-10-gained STAT3-bound sites at +1.5 kb and +11.5 kb of the Rgs3 TSS and +72.7 kb of the Rgs2 TSS. (F) RGS2 expression versus human metabolic and adipose-functional traits: HOMA-IR, WHR, waist, hip, BMI; circulating insulin, glucose, LDL, cholesterol, CRP, HbA1c, HDL; fat-cell volume and lipolysis readouts. (G) RGS3 siRNA versus scrambled siRNA knockdown validation. (H) pHSL western blots under RGS3 siRNA versus Src siRNA across the four conditions; pink arrowheads highlight loss of IL-10–mediated suppression in siRNA Rgs3 cells. (I) IL10-induced log2-fold-change heatmap of RGS3 protein normalized (Fig. 4B) to Tubulin across Ctrl and IL-10 conditions (diverging scale, Ctrl mean reference). (J) RGS3 western blot in lenti-RGS3 shRNA vs controls with RGS3 quantification (P = 0.0002). (K) RGS3 western blot in CRISPR RGS3KO vs controls with RGS3 quantification (P = 0.0001). (L) Quantification of IL10-induced RGS2 western blot (p=0.0037) (Fig. 4G). (M) RGS2 western blot in TregIL10KO versus TregIL10F/F WAT. (N) RGS2 levels in RGS2 siRNA versus scrambled siRNA knockdown validation and quantification (p<0.0005). (O and P) PKA-substrate western blots quantification in TregIL10KO versus TregIL10F/F (P = 0.0037) and iAdIL10RKO versus iAdIL10R^F/F (P = 0.0229) WAT. Mean ± SEM; statistics by two-tailed Student’s t-test or two-way ANOVA as appropriate.

