## Supplementary Tables for "Regulatory T cells establish an IL-10–IL10Rα immunometabolic checkpoint that limits HSL activation and lipolysis"

**Supplementary Table 1: Antobodies**

| Antibody | Company | Catalog # | Application (Dilution) |
| --- | --- | --- | --- |
| Anti-α-Tubulin | MilliporeSigma | CP06-100UG | WB (1:1000) |
| IRDye® 800CW Goat anti-Rabbit IgG Secondary Antibody | LICORBio | 926-32211 | WB (1:10000) |
| IRDye® 680RD Goat anti-Mouse IgG Secondary Antibody | LICORBio | 926-68070 | WB (1:10000) |
| IL13RA1 | Thermo Fisher Scientific | PA5-28309 | WB (1:1000) |
| KLF2 (E7K8Y) | Cell Signaling Technology | 51221 | WB (1:1000) |
| PTGFR | Thermo Fisher Scientific | PA5-34128 | WB (1:1000) |
| RGS3 | Thermo Fisher Scientific | PA5-79925 | WB (1:1000) |
| RGS2 | Proteintech | 10678-1-AP | WB (1:1000) |
| Anti-GAPDH | Thermo Fisher Scientific | PA1-9046 | WB (1:1000) |
| Phospho-HSL (Ser563) | Cell Signaling Technology | 4139 | WB (1:1000) |
| HSL | Cell Signaling Technology | 4107 | WB (1:1000) |
| il10ra | Proteintech | 13356-I-AP | WB(1:1000) |

**Supplementary Table 2: Cell surface markers used for flow cytometry.**

| **Blood, iWAT and eWAT** | |
| --- | --- |
| **Cell population** | **Cell surface markers*** |
| Leukocytes | CD45^+^ |
| Myeloid | CD45^+^CD11b^+^ |
| B cells | CD45^+^CD11b^-^CD90.2^-^B220^+^ |
| CD4 T cells | CD45^+^CD11b^-^CD90.2+CD3+ CD4^+^CD8b^-^ |
| CD8 T cells | CD45^+^CD11b^-^CD90.2+CD3+ CD4^-^CD8b+ |
| Regulatory T cells | CD45^+^CD11b^-^CD90.2+CD3+CD4^+^CD8b^-^CD25^+^FoxP3^+^ |
| IL10^+^ Regulatory T cells | CD45^+^CD11b^-^CD90.2+CD3+CD4^+^CD8b^-^CD25^+^FoxP3^+^IL10^+^ |
